# Densely sampled stimulus-response map of human cortex with single pulse TMS-EEG and its relation to whole brain neuroimaging measures

**DOI:** 10.1101/2024.06.16.599236

**Authors:** Yinming Sun, Molly V. Lucas, Christopher C. Cline, Matthew C. Menezes, Sanggyun Kim, Faizan S. Badami, Manjari Narayan, Wei Wu, Zafiris J. Daskalakis, Amit Etkin, Manish Saggar

**Affiliations:** Department of Psychiatry and Behavioral Sciences, Stanford University, Stanford, CA, USA; Department of Psychiatry, University of California San Diego, San Diego, CA, USA; Alto Neuroscience, Los Altos, CA

**Keywords:** TMS-EEG, community detection, multimodal, DWI, resting state fMRI, resting state EEG

## Abstract

Large-scale networks underpin brain functions. How such networks respond to focal stimulation can help decipher complex brain processes and optimize brain stimulation treatments. To map such stimulation-response patterns across the brain non-invasively, we recorded concurrent EEG responses from single-pulse transcranial magnetic stimulation (i.e., TMS-EEG) from over 100 cortical regions with two orthogonal coil orientations from one densely-sampled individual. We also acquired Human Connectome Project (HCP)-styled diffusion imaging scans (six), resting-state functional Magnetic Resonance Imaging (fMRI) scans (120 mins), resting-state EEG scans (108 mins), and structural MR scans (T1- and T2-weighted). Using the TMS-EEG data, we applied network science-based community detection to reveal insights about the brain’s causal-functional organization from both a stimulation and recording perspective. We also computed structural and functional maps and the electric field of each TMS stimulation condition. Altogether, we hope the release of this densely sampled (n=1) dataset will be a uniquely valuable resource for both basic and clinical neuroscience research.

## Introduction

The brain accomplishes complex actions through intricate sequences of coordinated neuronal activity. To understand how such processes arise, researchers have studied the brain’s wiring by estimating structural and resting state-based connectomes (Honey et al., 2009; Van Essen et al., 2013; Yeo et al., 2011). However, it is unclear how the brain’s response to activation arises from such connections. We can directly stimulate brain regions across the cortex to understand location-specific activation and relate the evoked response to structural and resting state properties. While invasive stimulation experiments in humans (Keller et al., 2014; Penfield & Jasper, 1954; Waters et al., 2018) and other species (Deisseroth et al., 2015) help address this gap, non-invasive approaches are necessary for understanding human neurocircuitry and providing a bridge to clinical applications.

Here, we used concurrent single-pulse transcranial magnetic stimulation and electroencephalography (TMS-EEG) to non-invasively map stimulus-response properties across the accessible human neocortex. Most previous TMS studies have targeted the motor cortex or dorsolateral prefrontal cortex (Tremblay et al., 2019). Even with one or a few targets, quantifying TMS evoked potentials (TEPs) effectively differentiates brain regions (Rosanova et al., 2009), tracks brain states (Massimini et al., 2005), and distinguishes patient populations from healthy controls (Ferrarelli et al., 2008; Voineskos et al., 2019). Rarely, however, have TEPs been used to map stimulus-response relationships across the broader cortical anatomy, partly due to the labor-intensive nature of TMS-EEG studies. The most highly-sampled study to date involved stimulating 18 target locations across the cortex in healthy subjects (Harquel et al., 2016), which is still relatively sparse and incomplete representation of cortical topography, especially relative to whole-brain imaging tools such as MRI. Moreover, while TMS-EEG has been used to assess coil orientation sensitivity (COS) of specific regions (Bonato et al., 2006; Casula et al., 2022), COS may differ across stimulation locations, which could be, in turn, informative for positioning the TMS coil in a previously unexplored brain region.

Therefore, to densely map stimulation responses as a function of location, we assessed cortical responsivity using TMS-EEG across the entirety of the accessible cortex, resulting in over 100 stimulation locations in one healthy individual (**Figure 1a-b**). We assessed the response from two orthogonal coil orientations for each stimulation location with a figure-8 coil. To anchor the TMS-EEG mapping results with more established measures from large-scale neuroimaging studies, we acquired Human Connectome Project (HCP)-styled diffusion imaging scans (six), resting-state functional Magnetic Resonance Imaging (fMRI) scans (120 mins), resting-state EEG scans (108 mins), and structural MR scans (T1- and T2-weighted). We generated standard whole-brain maps of structure and function from the high-fidelity anchoring measures, including myelination, cortical thickness, structural connectivity, functional connectivity, and EEG-based power envelop connectivity (Hipp et al., 2012; Toll et al., 2020) (**Figure 1i-m**).

**Figure 1.**
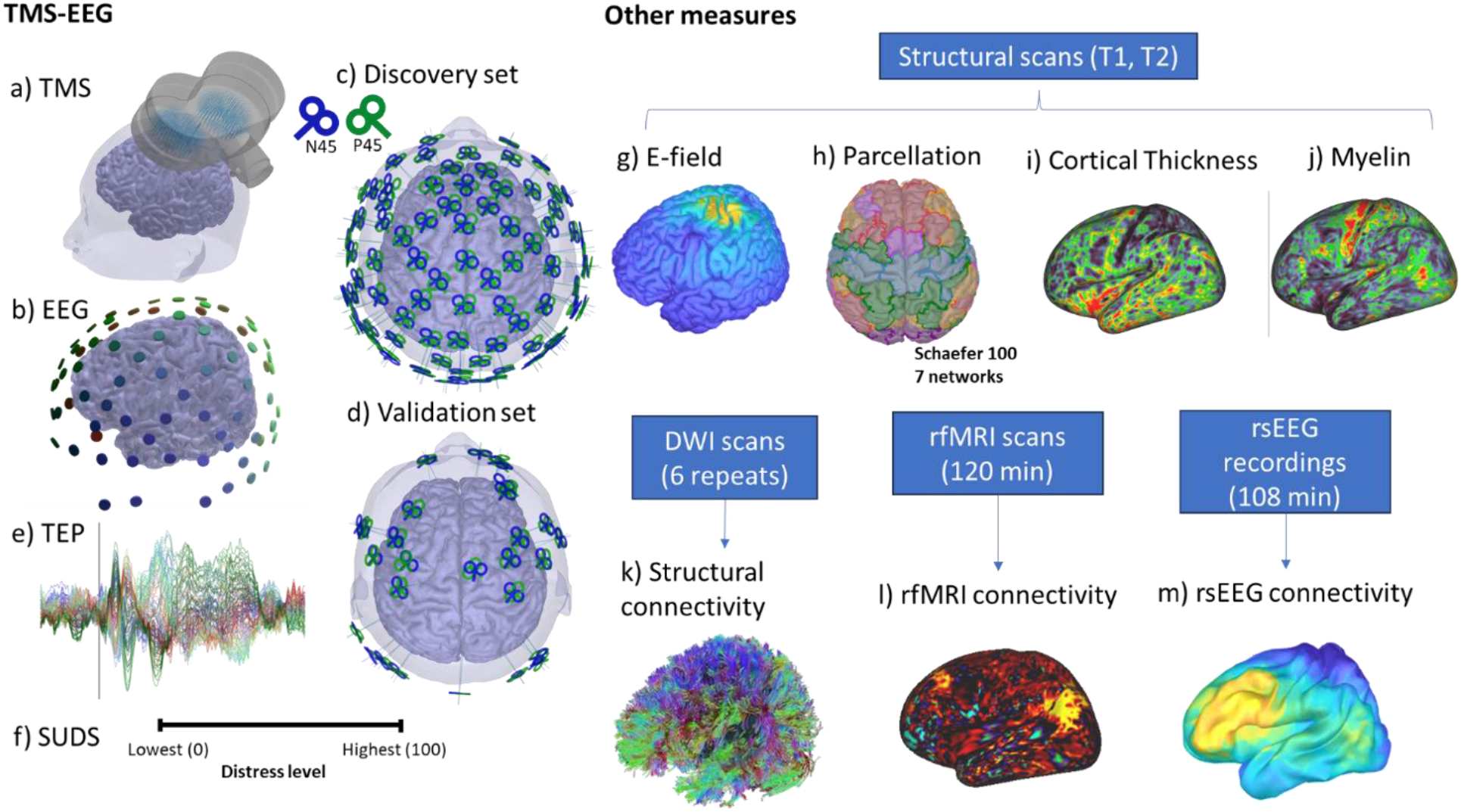
Experimental setup and components of the dataset. a) TMS stimulation was applied with a figure-8 coil, and b) EEG was recorded with a 95-channel system. c) Targets comprised a discovery set with 95 locations across the scalp based on the 10-5 international system and d) a validation set with 27 cortical locations selected to be roughly spaced across fMRI connectivity-based brain networks. Each location was stimulated twice with coil orientations approximately 90 degrees apart. e) The analyses focused on time points between 20 ms and 500 ms after each TMS stimulus, which were averaged across the trials (100 total) to produce the TMS Evoked Potential (TEP). f) Subjective units of distress scale (SUDS) values were also collected after each stimulation condition. In addition to the TMS-EEG measurements, we also collected T1 and T2 structural scans, which were used to compute g) stimulation E-field for each condition, h) parcellation, i) cortical thickness, and j) myelin. We also collected 6 diffusion scans based on the human connectome project (HCP) to generate the k) tractography and structural connectivity, l)120 min of resting fMRI to generate the resting state functional connectivity, and m) 108 min of resting EEG to generate power envelop connectivity for 4 typical EEG frequency bands (theta, alpha, beta, gamma).

To provide a quantitative overview of the TMS-EEG data, we applied network science-based community detection (Rubinov & Sporns, 2011) to reveal insights about the brain’s causal functional organization from both a stimulation and recording perspective. We also characterized how each stimulation-based community differed regarding structural and resting state measures. We hypothesized that inherent patterns of responses may exist across stimulation conditions based on measures of their TEP waveforms. We also hypothesized different brain regions have different levels of COS, with sensorimotor regions having high and associative cortical regions having low COS. Finally, we hypothesized that the patterns identified by community detection would differ significantly for specific structural and resting state measures.

We hope the release of this unique dataset (raw and derived), along with our preprocessing and analysis approaches, will guide basic researchers and clinicians alike in hypothesis generation and experimental planning involving TMS stimulation.

## Methods

### Experimental design

This study aimed to record TMS-EEG data from a highly dense grid of cortical targets to understand how spatial location impacts TMS-evoked response within an individual.

This study was approved by the Institutional Review Boards at Stanford University and the Palo Alto division of the Veteran Affairs Health Care System. One healthy, right-handed male (31 years old at study onset) voluntarily participated (author Y.S.). The participant completed an informed consent process before enrollment in the study.

The participant completed 18 TMS-EEG recording sessions over 2 months (each scheduled at the same time of day for consistency). Resting state EEG (rsEEG) was recorded (3 min eyes open, 3 min eyes closed) to measure baseline brain state (18 sessions). Following this, the participant completed between 6 and 22 conditions of TMS-EEG recording, as time allowed, in a predetermined randomized order. In total, 244 stimulation conditions were completed. Following each condition, the participant rated the level of discomfort on a Subjective Units of Distress scale (SUDs) ranging from 0 to 100 (100 being maximal discomfort). Each condition or target (location x coil angle) contained 100 TMS pulses with an inter-stimulus interval (ISI) of 2 seconds (jittered: +/- 200 ms).

To anchor the TMS-EEG mapping results with more established measures from large-scale neuroimaging studies, the participant also completed 6 diffusion scans based on the human connectome project (HCP) (Glasser et al., 2013; Sotiropoulos et al., 2013), 120 min of resting fMRI, and 108 min of resting EEG, along with T1 and T2 scans. All MRI data was collected at the Stanford Center for Cognitive and Neurobiological Imaging (CNI) using a General Electric (GE) Discovery MR750 3T scanner and a 32-channel Nova head coil. Scan sessions were scheduled for approximately the same time of day in the afternoon.

### TMS-EEG stimulation targets

A structural MRI scan was used to define TMS stimulation targets in native space and allow for real-time neuronavigation for TMS targeting. The T1-weighted structural scan was acquired using a GE 3D BRAVO sequence (sagittal, 256 slices, 1 mm isotropic resolution, echo time (TE) = 3.44 ms, repetition time (TR) = 8.69 ms, inversion time (TI) = 500 ms, flip angle (FA) = 11 degrees) at the Stanford Center for Cognitive and Neurobiological Imaging (CNI) using a General Electric (GE) Discovery MR750 3T scanner and a 32 channel Nova head coil.

TMS target locations (N = 122) were pre-defined using the subject’s native space 1 mm T1 structural scan. Of these 122 target locations, the main dataset comprised 95 cortical locations based on the 96-channel cap electrode positions (based on the 10-5 international system). The validation dataset comprised 27 cortical locations selected to be roughly spaced across fMRI connectivity-based brain networks. The validation locations were interspersed between the electrode-based targets.

The electrode-based (i.e., main) locations were defined by projecting the 10-5 electrode positions to the native-space brain mesh (Giacometti et al., 2014). The fMRI brain network locations were created by converting template-space coordinates into the subject’s native space using an automated algorithm developed within the lab based on a previous population study (Toll, 2018; Toll et al., 2020).

Each location was targeted twice using two coil orientations. The first orientation (N45) oriented the coil handle towards the back of the brain and 45 degrees clockwise from the anterior-posterior axis. The second orientation (P45) rotated the coil handle 45 degrees counterclockwise. In total, this created 244 stimulation conditions.

### TMS-EEG data acquisition

TMS-EEG data was recorded using a 96-channel Brain Products actiCAP slim cap and actiCHamp amplifier at 25 kHz sampling rate. TMS stimulation was applied using a MagVenture B65 coil and a MagVenture R30 stimulator.

### Stimulation intensity

The intensity was set at 120% of the resting motor threshold (RMT). RMT was the stimulation amplitude that elicited > 50 µV motor evoked potential (MEP) in 5 out of 10 consecutive trials. MEPs were recorded using a Biopac System, where the active electrode was placed on the left first dorsal interosseous (FDI) muscle. Using this method, the stimulator output peak dI/dt corresponded to 134 A/µs at 120% of RMT.

### Auditory masking

Auditory masking was used throughout the experiment in the form of Bose noise-cancelling headphones playing white noise. White noise volume was set each day based on participant feedback; volume was increased until either the participant could not hear the TMS firing or the volume became uncomfortable for the participant.

### Neuronavigation

Brainsight neuronavigation software (Rogue Research) was used to position and track each stimulation target (i.e., location and angle) throughout the experiment. Single-trial level targeting information was recorded in Brainsight, measuring 6 degrees of freedom.

### MRI data acquisition

#### Structural MRI

A T_1_-weighted structural scan was acquired using a GE 3D BRAVO sequence (sagittal, 256 slices, 1 mm isotropic resolution, echo time (TE) = 3.44 ms, repetition time (TR) = 8.69 ms, inversion time (TI) = 500 ms, flip angle (FA) = 11 degrees). This structural scan was used to define stimulation targets and for in-session neuronavigation.

For electric field modeling, one T_1_-weighted and one T_2_-weighted scan were acquired with 0.5 mm x 0.5 mm x 0.8 mm voxel resolution (T_1_-weighted scan parameters: sagittal, 232 slices, TE = 3.58 ms, TR = 7.884 ms, TI = 1100 ms, FA = 9 degrees; T_2_-weighted scan parameters: sagittal, 228 slices, TE = 96.96 ms, TR = 2502 ms, FA = 90 degrees).

### Diffusion-weighted imaging

Diffusion-weighted imaging (DWI) scans were acquired with a GE-adapted multiband sequence based on the Human Connectome Project (Glasser et al., 2013; Sotiropoulos et al., 2013). Six identical sets of scans were acquired over three days. Four separate scans were acquired for each set, pieced together during processing into one scan with 150 directions distributed across 2 shells with gradient field values of 1500 and 3000 s·mm^-2^ (1.5 x 1.5 x 1.5 mm voxels resolution). The four scans consisted of a pair of scans for each shell in opposite phase encoding directions (anterior to posterior and posterior to anterior).

For the purpose of quantifying white matter conductivity in electric field modeling of the TMS stimulus, an additional 60-direction DWI scan was acquired in an earlier session in accordance with the recommendations of the SimNIBS preprocessing pipeline (Opitz et al., 2011). The slice thickness was 2.1 mm.

### Resting-state fMRI

Twelve 10 min resting-state fMRI (rsfMRI) scans with a GE gradient echo planar imaging (EPI) sequence (oblique slices, 2.9 mm slice thickness, TR = 2000 ms, TE = 30.0 ms, FA = 77 degrees) were collected over the course of 4 sessions (120 min in total) with gradient field maps for each session. During each scan, the participant was in the supine position with eyes focused on a fixation cross displayed overhead.

### MRI data processing

Neuroimaging data (T1, T2, DWI, rsfMRI) were preprocessed with the minimal Human Connectome Project (HCP) preprocessing pipeline (Glasser et al., 2013). Since details of the pipeline have been extensively presented by the HCP group, only relevant experimental specific configurations are mentioned here.

The 0.8 mm T1 and T2 scans were used for the structural portion of the pipeline since the increased resolution allows for a more accurate estimation of structural measures, especially myelination. As recommended by the HCP pipeline, DWI scans from the same day were combined before applying the HCP diffusion pipeline. All 6 sets of DWI scans were co-registered to the same T1, concatenated across volumes, and masked within the brain to optimize the subsequent white matter tract reconstruction. Each rsfMRI scan was corrected for distortions with the corresponding GE B0 fieldmap.

### Computing rsfMRI dense connectome

A Connectivity Informatics Technology Initiative (CIFTI) dense connectome was calculated for each preprocessed rsfMRI dense time series using the -cifti-correlation command from connectome workbench. To avoid the effect of transients at the start of the scan, 10 s (or 5 timepoints) of data from the beginning of the rsfMRI time series was removed before computing the correlation. The CIFTI dense connectome matrices from all 12 scans were imported into Matlab and averaged to produce a more reliable estimate of the dense connectome for further processing.

### DWI reconstruction and fiber tracking

The 6 DWI scans was co-registered to the T1 structural scan output from Freesurfer as part of the SimNIBS processing pipeline and then concatenated. With 6 scans combined, the total number of diffusion sampling directions were 450 for each of the two shells. Diffusion reconstruction and fiber tracking was done in DSI studio with the combined scan (Yeh et al., 2013). The diffusion data were reconstructed using generalized q-sampling imaging (GQI) (Yeh et al., 2010) with a diffusion sampling length ratio of 1.25. The GQI method was used since it can accurately capture crossing fibers and was shown to produce the most accurate results in combination with deterministic tractography based on recent literature comparing different reconstruction methods (Maier-Hein et al., 2017). For deterministic fiber tracking, the quantitative anisotropy threshold was 2.24. The angular threshold was randomly selected from 15 degrees to 90 degrees. The step size was randomly selected from 0.5 voxel to 1.5 voxels. The fiber trajectories were smoothed by averaging the propagation direction with a percentage of the previous direction. The percentage was randomly selected from 0% to 95%. Tracks with length shorter than 30 mm or longer than 300 mm were discarded. A total of 100000 tracts were calculated.

### rsEEG power envelope connectivity

For rsEEG data, the native cortical mesh was downsampled to 3000 vertices from the 15002 vertices mesh and the headmodel was calculated based on the channel position from session 6 (a randomly chosen and representative session). The co-registered atlases were also downsampled accordingly. wMNE source estimation was completed in accordance with prior rsEEG studies from the lab (Toll et al., 2020).

rsEEG power envelope connectivity (PEC) between the vertices of the downsampled cortical mesh was calculated in accordance to previously published methods (Hipp et al., 2012; Toll et al., 2020). For each session, a separate connectivity matrix was calculated for each condition (i.e., eyes open and eyes closed) and for the standard frequency bands of theta (5 – 8 Hz), alpha (8 – 12 Hz), beta (13 – 24 Hz), and gamma (25 – 50 Hz). Each connectivity matrix was then normalized based on its mean and standard deviation across all connections. Finally, the normalized connectivity matrices were averaged across sessions, resulting in 8 average connectivity matrices (2 conditions x 4 frequency bands).

### Simulating TMS induced electric field

Realistic electric field modeling was completed using SimNIBS based on individual anatomy (T1, T2, and DWI) (Opitz et al., 2011; Saturnino et al., 2019), coil modeling based on the MagVenture Cool-B65 coil, and precise stimulation targets collected from Brainsight during experiment.

A dipole model of the coil was created in accordance to the method described by Thielscher and colleagues (Thielscher & Kammer, 2002). The dimensions of the coil were based on manufacturer’s specifications for the Magventure B65 and verified by X-ray imaging. The figure-8 coil consisted of two flat circular coils in the same plane, each with an inner radius of 17.5 mm and an outer radius of 37.5 mm. Each circular coil had 2 layers of 5 windings with each layer of winding having a height of 6 mm.

The T1 and T2 MRIs used were the 0.8 mm voxel scans, while the DWI scan used was a 60-dir single shell scan. The set of scans were collected in accordance to that suggested for SimNIBS at the time of the experiment. The direct simulation output of SimNIBS is a volumetric mesh bounded by native surfaces. To enable various further processing, the output results were also mapped onto Native and Fsaverage cortical surfaces, as well as native and MNI warped NIfTI volumes.

To obtain the stimulation target specifications for SimNIBS simulation, the recorded Brainsight targets were converted to the format expected for SimNIBS. Brainsight outputs the stimulation targets in world coordinates along with a set of three vectors, which points in the posterior to anterior, left to right, and ventral to dorsal directions of the figure-8 coil. Since a different T1 was used for targeting before a higher resolution T1 was obtained for simulation, an affine transform was calculated for transforming the target T1 used for Brainsight targeting to the T1 used for SimNIBS simulation. The resulting transform was applied to the stimulation target locations outputted by Brainsight.

### Computing stimulation ROI-based measures

Since neuronal activation depends on the induced electric field’s strength, the cortex region activated by a TMS pulse can be approximated by vertices in the cortical mesh with E-field values exceeding a certain activation threshold. Given previous literature on the range of TMS E-field values(Rosanova et al., 2009) and ensuring each target has at least a few vertices exceeding the threshold, stimulation regions of interest (ROIs) were defined with an E-field threshold exceeding 70 V/m. The size of the stimulation ROI and the associated average intensity were computed for the 15002 vertices native cortical surface used for source localization of TMS-EEG signals.

### Myelination and cortical thickness

Stimulation ROI values were extracted in the Fsaverage surface space for myelination and cortical thickness since the E-field maps have already been converted to this space as part of the SimNIBS simulation. Specifically, the myelination and thickness values in the native fs_LR space were warped to the Fsaverage space using Connectome Workbench commands. Once in the same space, values from the vertices part of each ROI were averaged, producing 190 ROI values (95 targets x 2 orientations).

### Diffusion tractography

For diffusion tractography, stimulation ROI values were extracted in the T1 volume space since the tracts calculated from DSI Studio can only be seeded in volume space. Specifically, the grey matter region of the cortex in the reference T1 was extracted using the associated segmentation mask generated as part of SimNIBS. The resulting masked E-field volume was then subjected to the same E-field threshold of 70 V/m to extract the activated brain region as a volumetric ROI.

After the diffusion image was co-registered to the same T1 image, the volumetric stimulation ROIs were imported into DSI Studio. The Schaefer atlas (7-network, 100 parcellation version)(Schaefer et al., 2018) in MNI space was also imported into DSI Studio and warped to the T1 native space to obtain 100 response ROIs.

Connectivity between the 190 stimulation ROIs (95 channels x 2 orientation) and the 100 response ROIs were generated based on the tract count that begins in a stimulation ROI and end in a response ROI, resulting in a 190 by 100 non-symmetrical matrix. To control for the effect of ROI area, the tract counts were normalized by the geometrical average of the areas of the connected ROIs (Betzel et al., 2019).

### rsfMRI connectivity

For rsfMRI connectivity, the stimulation ROI values were extracted in the CIFTI space since the rsfMRI dense connectome was calculated in that space. The E-field based ROIs in FSaverage space were warped to the same space using Connectome Workbench commands. The connectivity matrix, the warped stimulation ROIs, along with 100 ROI definitions in the same CIFTI space were subsequently imported to MATLAB for further processing. The large connectivity matrix was imported with the CIFTI toolbox provided by HCP.

To calculate the connectivity between E-field-based ROIs and the 100 response ROIs, the values of the connectivity matrix belonging to rows and columns defined by the two sets of ROIs were averaged. The medial wall indices were removed from the E-field surface indices before the correct corresponding values were extracted from the CIFTI connectivity matrix. The result was a 190 by 100 ROI connectivity matrix.

### rsEEG connectivity

For rsEEG, since the vertex-based power envelope connectivity was calculated for a 3000 vertices native surface, the E-field maps in higher resolution native space were projected to the same surface with Brainstorm before applying the 70 V/m threshold to define the ROIs. Once the stimulation ROIs were defined in the same space as the rsEEG data, values of the vertex connectivity matrix belonging to rows and columns defined by the E-field and 100 response ROIs, respectively were averaged. Eight 190 by 100 ROI connectivity matrices were calculated, corresponding to the 8 vertex-based connectivity matrices.

### TMS-EEG data preprocessing

Continuous data was epoched (-1 to 1.5 seconds surrounding each pulse). The primary TMS artifact was removed (-2 to 12 ms) and the timespan was interpolated using autoregressive interpolation. Data was downsampled from 25 kHz to 1 kHz and baseline corrected. A modified high-pass filter (1 Hz) was applied to minimize the spread of high amplitude residual pulse artifact to surrounding pre- and post-pulse data. A data-driven Wiener noise estimate was calculated for each channel; bad channels were identified as those exceeding a threshold of 10. Source-estimate utilizing a noise-discarding algorithm (SOUND) was used to identify and remove noise from the data (lambda regularization parameter = 10^-1.5^) (Mutanen et al., 2018). Large secondary stimulation artifact was removed using a decay fitting process. Line noise was attenuated using a Butterworth bandstop filter (58 – 62 Hz). ICA was used to decompose scalp signal into independent components. ICLabel was used to label each component (Pion-Tonachini et al., 2019). Components that exceeded threshold (eye score > 0.08, muscle score > 0.2, or brain score < 0.3) were rejected. To identify time-locked muscle artifact components, the mean absolute component amplitude ratio was calculated, comparing the 11 to 30 ms time window to that of the whole epoch. If the ratio exceeded 8, the component was rejected. Finally, components with a peak amplitude greater than 15 uV were rejected, as this amplitude is inconsistent with biological activity. A low-pass filter (200 Hz) was applied. The data was average re-referenced. A detailed description of these methods is presented elsewhere (Cline et al., 2021). This pipeline consistently removed artifacts across the broad range of stimulation locations across the scalp.

### Expert rating of TEP signal quality

To ensure the signal quality of the TEPs while maintaining objectivity and replicability, the preprocessed TEPs were separately rated on an integer scale from 0 to 5, with 5 being the highest quality, by three raters with expertise in TMS-EEG analysis. Expert raters independently assigned a rating score based on only TEP features, such as amplitude, topography, and noise floor, with knowledge of the stimulation condition (location, orientation). The list of target prompts was randomized differently for each rater to prevent potential order effects in the rating process. The average score and the maximum score differences were calculated based on the scores of individual expert raters. Targets with scores that had a maximum score range of 2 or more were discussed and assigned a final score based on consensus. Targets with an average score greater than 2 and less than 3 were also discussed and assigned a final score based on consensus. Consensus scores were assigned with steps of 0.5. Average scores from conditions without discussion were floored to steps of 0.5 (e.g., 3.67 was floored to 3.5) and considered final. A cutoff score for an acceptable TEP was 2 or more for this study. This resulted in 198 conditions being used for subsequent analysis. Supplemental Fig. 1 indicates which target conditions were not included in subsequent analysis. Online tables show the TEP response from all target locations that passed quality checks (https://figshare.com/s/a39fa560d79b563f0f49).

### Community detection analysis

Community detection based on measures of modularity(Rubinov & Sporns, 2011) was applied to determine A) similarity and B) coil orientation sensitivity across 1) stimulation targets and 2) recording channels, which answered four complementary questions.

For each analysis, community detection was also applied to TEP based connectivity, which was calculated for each stimulation condition by correlating the channel TEP timeseries with each other, resulting in a 95 by 95 matrix. The channel TEP timeseries consist of time points between 20 ms and 500 ms post-stimulus (i.e., 481 data points).

With TEP timeseries data, each row of the input matrix is the timeseries data across a condition, which is either across all channels for a given stimulation condition or across all stimulation conditions for a given channel. With TEP connectivity data, the connectivity matrix elements were first linearized and combined across all available conditions before applying community detection.

Community detection was applied with the Louvain algorithm with asymmetric treatment of negative weights (Bertolero et al., 2017; Rubinov & Sporns, 2011). Community detection was run 1000 times to obtain the mean modularity score (𝑄) and consensus assignment. The modularity score was calculated based on equation 1.

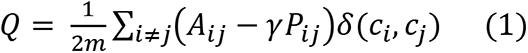

𝑄 is the modularity of a network with 𝑛 nodes and 𝑚 edges

Edge weight: 𝐴_𝑖𝑗_ = 𝑒𝑑𝑔𝑒 𝑤𝑒𝑖𝑔ℎ𝑡 𝑏𝑒𝑡𝑤𝑒𝑒𝑛 𝑖 𝑎𝑛𝑑 𝑗

Node strength: 𝑘_𝑖_ = ∑_𝑗_ 𝐴_𝑖𝑗_

Probability of connection between nodes 𝑖 and 𝑗 in random network: 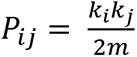

𝛾 is the resolution parameter and set to 1, 𝑐_𝑥_ is the community of node 𝑥

### Community detection across stimulation conditions

Supplemental Fig. 2 shows the steps involved in running community detection across stimulation conditions. For each stimulation target, the TEP data matrix (95 channels x 481 timepoints) was correlated across timepoints to obtain a 95 x 95 channel connectivity matrix, which was then reshaped into a 1 x 9025 vector and combined. The resulting targets by channel connectivity matrix was correlated across all connectivity values to produce a targets x targets matrix (𝐴_𝑖𝑗_ in equation 1) that was used as input for community detection. For the discovery set, matrix 𝐴_𝑖𝑗_ has dimensions of 143 x 9025, while for the validation set, matrix 𝐴_𝑖𝑗_ has dimensions of 42 x 9025.

The validation set was used to compare the community assignment based on 1) the nearest neighbor based on Euclidean distance, and 2) top 5 targets based on correlation between TEP connectivity (highest frequency community with highest average correlation).

Coil orientation sensitivity (COS) for each stimulation location was quantified by the community agreement between the associated P45 and N45 targets. If paired targets were assigned to different communities, then the shared location was considered to have high COS.

### Comparing structural and resting state measures across communities

With stimulation conditions assigned into different communities based on TEP measures, differences in structural and resting state measures across communities can be readily quantified. Specifically, we compared measures of cortical thickness, myelination, structural connectivity, rfMRI connectivity, and EEG based power envelop connectivity across different frequency bands. As specified in the earlier sections, target specific measures of cortical thickness and myelin was defined as the average value across the E-field defined ROI, which was set at 70 V/m. For the connectivity features (i.e., structural, rsfMRI, and rsEEG), we chose degree as the metric, which was defined as the sum of the measure between the E-field ROI and Schaefer 100 7-network parcels. A global metric was obtained by summing the connectivity measure across all the parcels, while network specific metrics were obtained by dividing the sum of the connectivity measure across only parcels of a specific network by the total sum across all parcels.

All measures were compared across the communities with the Kruskal-Wallis test after regressing out SUDs as a covariate. SUDs were considered a measure of the somatosensory confound associated with each stimulation condition. If there were a significant group effect, paired comparisons between the communities were performed with a rank-sum test and subsequent Tukey correction. Non-parametric tests were used to avoid assumptions of normality.

### Community detection across recording channels

Supplemental Fig. 3 shows the steps involved in running community detection across recording channels. For each recording channel, an associated targets x timepoints matrix can be correlated across timepoints to obtain a targets x targets similarity matrix, which can then be reshaped into a vector and combined. The resulting channels by targets connectivity matrix was used as the input (𝐴_𝑖𝑗_ in equation 1) for community detection. Since the number of targets for the discovery set and validation set are 143 and 42 respectively, the associated matrix 𝐴_𝑖𝑗_ has dimensions of 95 x 20449 and 95 x 1764 respectively.

COS for each channel 𝑗 was quantified by the modularity score of the TEP timeseries across all targets using coil orientation as the community label. Targets were split into orthogonal and parallel groups, which is in reference to the central sulcus for each hemisphere, before computing the correlation. A higher modularity score means a higher COS. A null distribution was generated by shuffling the community assignments and then computing the modularity score by chance 1000 times. A significance test (p < 0.05, or above 95% random values) was performed by comparing the actual modularity against that of the null distribution using exact statistics.

## QUANTIFICATION AND STATISTICAL ANALYSIS

All quantification and statistical tests were done in Matlab, which are described in different parts of the Methods Details section and collated below.

Across stimulation conditions, community detection was run 1000 times and averaged to produce a mean modularity and consensus assignment, which ensured the stability of the communities. Kruskal-Wallis tests were used to compare the structural and resting state measures across the stimulation-based communities. If there were a significant group effect, paired comparisons between the communities were performed with a rank-sum test and subsequent Tukey correction. Non-parametric tests were used to avoid assumptions of normality.

Community detection across recording channels was also run 1000 times and averaged to produce a mean modularity and consensus assignment. Since the same recording channels are used for the discovery and validation set conditions, channel community assignments determined from the discovery set can be directly applied to the validation set. A high modularity score for the validation set with similar connectivity patterns as the discovery set would suggest robust channel-based communities that are minimally dependent on the stimulation site.

To establish the significance of coil orientation sensitivity for each recording channel, a significance test (p < 0.05, or above 95% random values) was performed by comparing the actual modularity against that of the null distribution using exact statistics. The null distribution was generated by shuffling the coil orientation labels (i.e., the true community assignment) and then computing the modularity score by chance 1000 times. A significance test (p < 0.05, or above 95% random values) was performed by comparing the actual modularity against that of the null distribution using exact statistics.

## Results

### Summary of the multimodal dataset

Figure 1 illustrates the experimental setup and components of the dataset. To non-invasively map stimulation-response patterns across the brain, we recorded concurrent EEG responses from single-pulse transcranial magnetic stimulation (i.e., TMS-EEG) of over 100 cortical regions with two orthogonal coil orientations from one individual across 18 sessions. TMS stimulation was applied with a figure-8 coil, and EEG was recorded with a 95-channel system. The targets comprised a *discovery set* with 95 locations across the scalp based on the 10-5 international system(Oostenveld & Praamstra, 2001) and a *validation set* with 27 cortical locations selected to be roughly spaced across fMRI connectivity-based brain networks(Toll et al., 2020). In addition to using real-time coil tracking and state-of-art noise reduction techniques (e.g., auditory masking, coil padding), we also quantified the relative discomfort of each stimulation condition using the subjective units of distress scale (SUDS) as a potential signal regressor. To better remove the unique noise signatures of the never tested stimulation locations, we also designed a new preprocessing pipeline(Cline et al., 2021) and created a practical consensus rating system for rejecting noisy data after preprocessing (Supplemental **Fig. S1**). The analyses focused on time points between 20 ms and 500 ms after each TMS stimulus, which were averaged across the trials (100 total) to produce the TMS Evoked Potential (TEP).

In addition to the TMS-EEG measurements, we collected T1- and T2-weighted structural scans, which were used for simulating the electric field (E-field) for each condition and generating the brain parcellation (Schaefer 100, 7 networks(Schaefer et al., 2018)) for subsequent analysis. The structural scans were also used for computing cortical thickness maps and myelin(Glasser et al., 2013). The HCP-styled diffusion-weighted imaging was used to generate high fidelity structural connectivity from the whole brain tractography(Sotiropoulos et al., 2013) (Figure 1k); 120 min of resting fMRI was used to generate the resting state functional connectivity(Laumann et al., 2015) (Figure 1l); and 108 min of resting EEG was used to generate power envelop connectivity for 4 typical EEG frequency bands (theta, alpha, beta, gamma)(Hipp et al., 2012; Toll et al., 2020). Target-specific measures of cortical thickness and myelin map were defined as the average value across the E-field defined region of interest (ROI), which was threshold at 70 V/m, while the connectivity measures (i.e., DWI, rsfMRI, and rsEEG based) were defined from the E-field ROI to the rest of the brain across all defined parcellations.

### A look-up table of TMS-EEG responses

In addition to providing the raw and preprocessed data, an online lookup table was created for researchers to check the response profile of each stimulation condition easily. **Table 1** is a primer for the complete online tables found at 10.6084/m9.figshare.14701848 (https://figshare.com/s/a39fa560d79b563f0f49). Each row of **Table 1** reflects one stimulation condition. "Site" describes the center of the coil placement, and "Angle" describes the coil handle orientation. P45 has the coil handle oriented towards the back of the brain and 45 degrees counterclockwise from the anterior-posterior axis, while N45 has the coil handle oriented 45 degrees in the clockwise direction. Target coordinates are in the MNI space. The E-field map is generated by SimNIBS (coil not to scale). Sensor-space butterfly plots, topoplots, and source-space brain plots show the trial-averaged response to stimulation. GMFA shows the global mean field amplitude.

**Table 1:**
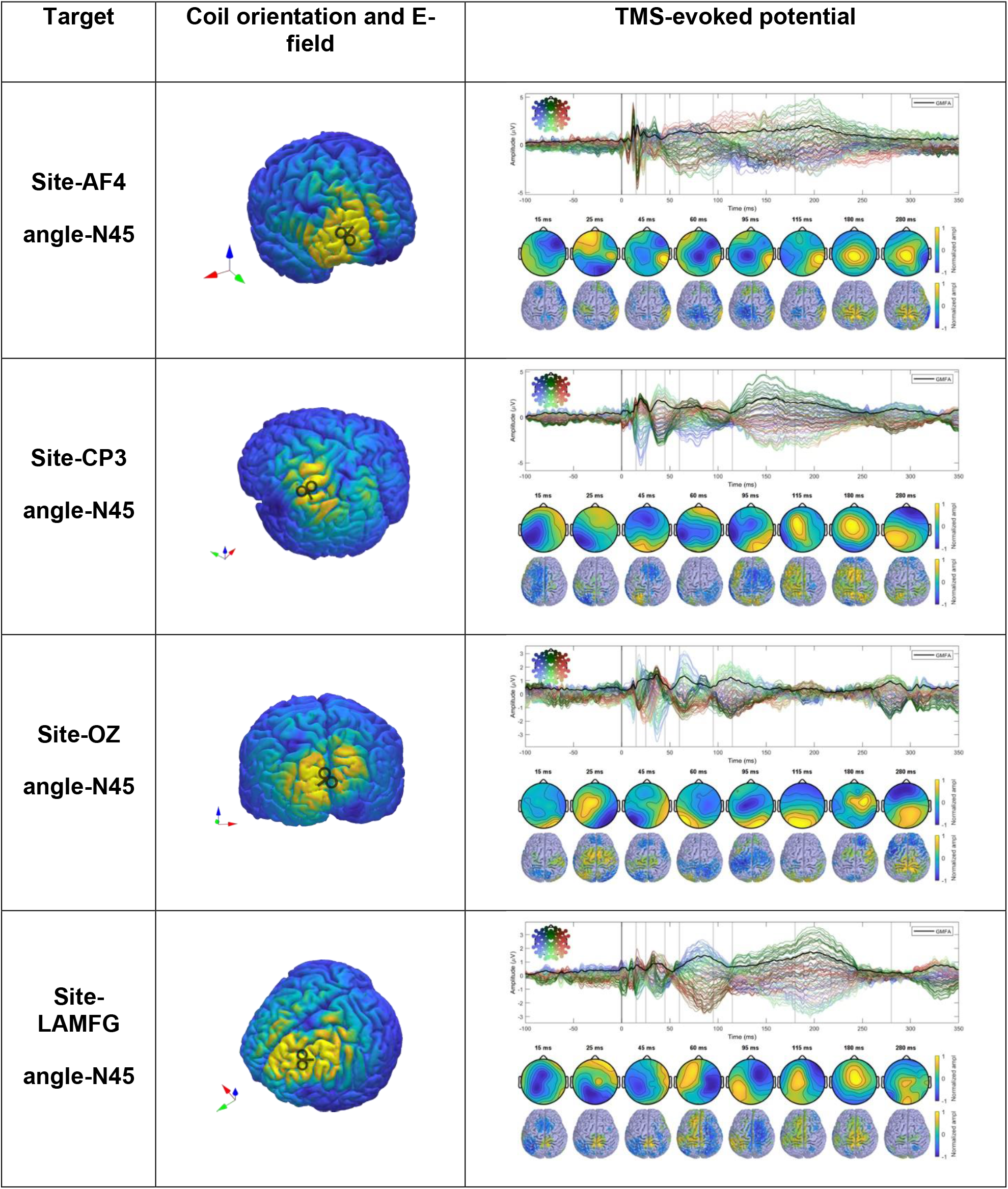
A look-up table of TMS-EEG responses.

### Clustering responses (TEPs) across stimulation conditions

To identify prominent patterns (or clusters of similar responses) across all the recorded TMS-EEG conditions, we applied network science-based community detection approaches, which are highly effective in capturing significant patterns in high-dimensional neuroimaging data(Rubinov & Sporns, 2011). The same approach was used to answer several complementary questions, as detailed below. To reduce any confounding effect of somatosensory sensation, we regressed the SUDS values across conditions from the TEP timeseries before running subsequent analyses.

The foremost question was how the stimulation conditions are related to each other. The answer to this question can help us understand the brain’s causal functional organization and target selection in treatments. For this analysis, we applied community detection to a TEP connectivity similarity matrix, which was derived by calculating the correlation between channel TEPs for each stimulation condition and correlating the resulting channel connectivity matrix elements. Further details are provided in the methods section (Supplemental **Fig. S2**).

Across all stimulation conditions, including both orientations, community detection with TEP connectivity similarity identified three communities in the discovery set (Figure 2a). Rather than being separated regionally, the targets were interspersed across cortical regions (Figure 2b). The assigned communities of the validation set also showed similar distributions. Comparing channel TEPs for each community showed both temporal and spatial differences. Figure 2c shows the absolute value channel TEPs averaged across targets of the same community with the topography shown for one selected peak. Supplemental **Figure S4** shows the same community TEPs plotted at their channel locations. From the butterfly plots in Figure 2c, we can see how different peaks (representing both positive and negative peaks of the TEP) are part of each community. Most notably, community 1 had the largest peak at around 43 ms, which was prominent in the lateral regions of both hemispheres (e.g., C3/4/5/6, CP3/4/5/6, P3/4/5/6). Community 2 had the largest peak at 191 ms, most prominent in the central midline regions (e.g., FCz, C1/2, CCP1h/2h, CP1/z/2, Pz). Community 3 had two large early peaks at 39 ms and 70 ms, the latter was most prominent in the frontal central and occipital regions (e.g., AFz, AFF1h/2h, F1/z/2, Oz, OI1h/2h). Community 2 had the latest noticeable peak at 316 ms, while community 3 had the greatest number of unique peaks among the communities.

**Figure 2.**
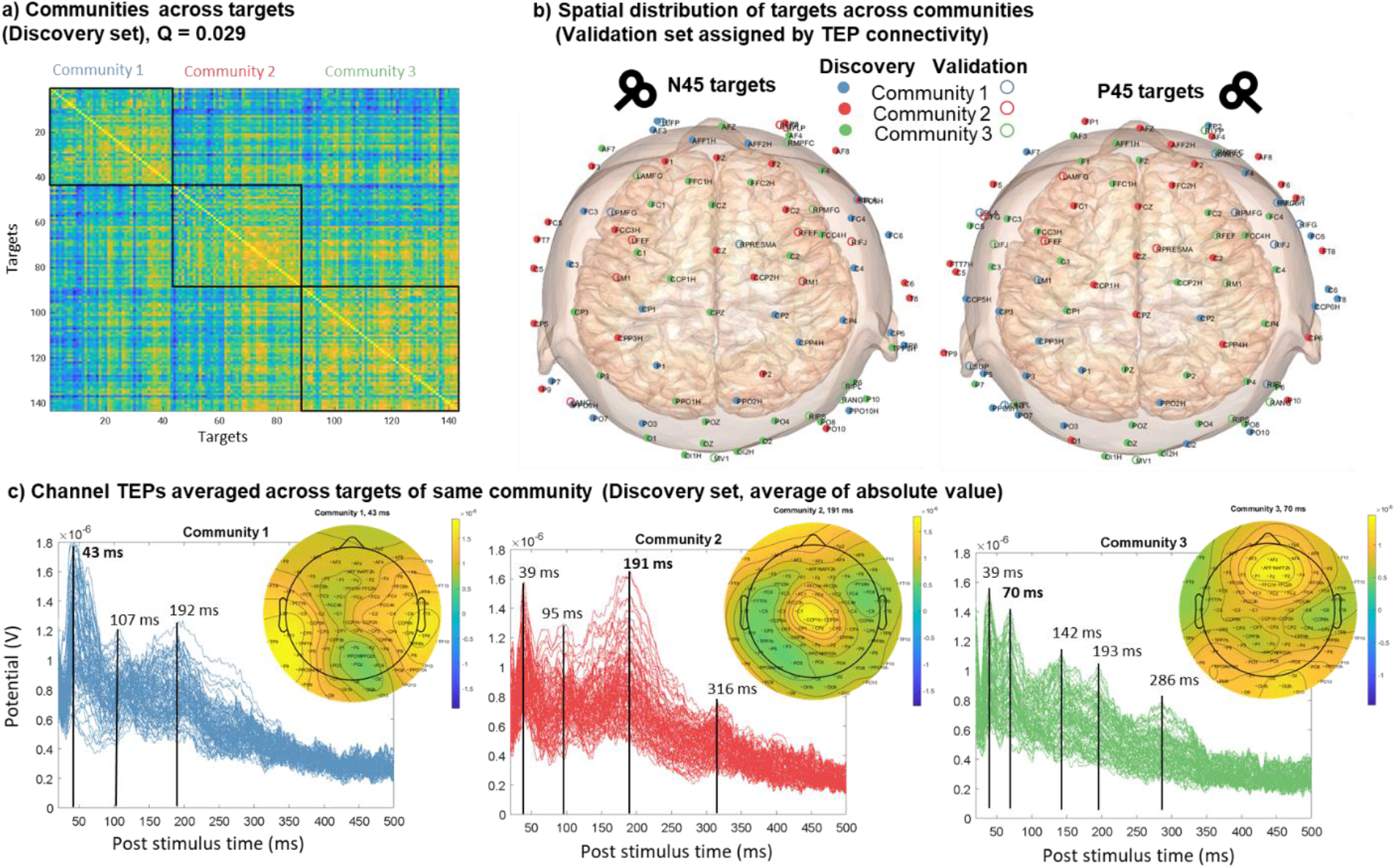
Clustering responses (TEPs) across stimulation conditions. a) TEP connectivity similarity matrix showing three communities in the discovery set. b) The targets are interspersed across cortical regions for the discovery set. The assigned communities of the validation set also show similar distributions. c) Channel TEPs averaged across targets of same community (Discovery set, average of absolute value) for each of the communities. The inset topoplots show the amplitude distribution at 43 ms, 191 ms, and 70 ms for community 1, 2, and 3 respectively.

### Coil orientation sensitivity across stimulation locations

Since previous studies with the figure-8 coil have shown orientation sensitivity, especially for the primary motor cortex(Bonato et al., 2006), we examined how sensitive each stimulation location is to coil orientation. For this analysis, we built upon our first community-detection-based clustering analysis by checking the agreement of the community assignments across two orientations. Thus, if the two orientations at the same site share the same community, then that site can be considered to have low coil orientation sensitivity (COS). Conversely, the site is considered to have high COS if the two orientations differ.

Our results showed that except for targets over the occipital region (e.g., Oz, PO3/z/4, OI1h/2h, MV1), target orientations of the same site do not share the same community, meaning they have high COS (Figure 3a). The same was true for the validation targets based on the agreement between assigned values. The percentage of low/high COS locations is similar for the discovery and validation sets, which are 37.5/62.5 and 33.9/66.1, respectively.

**Figure 3.**
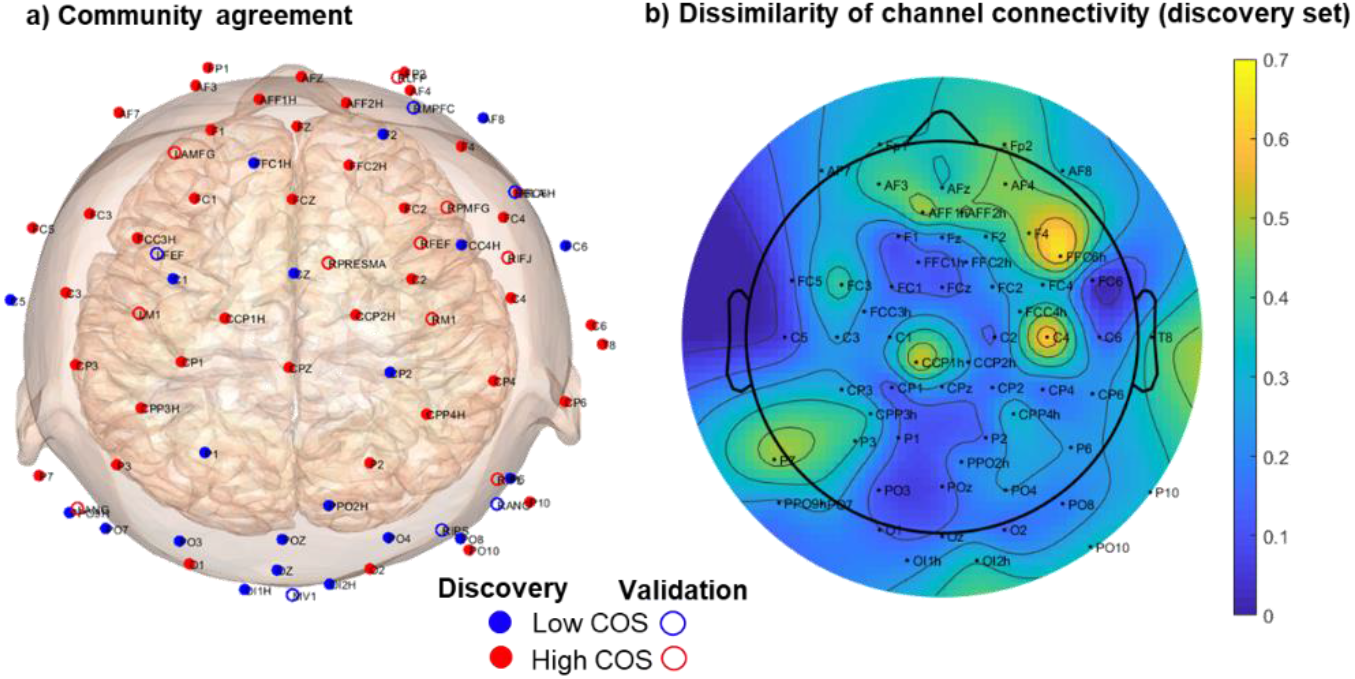
a) Community matching between the two orientations shows that except for the posterior central and occipital regions, most paired targets are not part of the same community. b) Dissimilarity between the channel connectivity of paired conditions for the discovery set shows a similar pattern for COS, but with continuous values.

In addition to our binary quantification of COS, we also checked the dissimilarity between the channel connectivity of paired conditions for the discovery set as a more continuous measure of COS. We defined dissimilarity as 1 minus Pearson’s correlation between paired orientations, meaning higher dissimilarity translates to higher COS. Similar to our findings with community agreement, midline posterior, and occipital electrodes had the lowest COS. At the same time, in line with previous results, the results show the highest relative COS to be over the right motor cortex (i.e., C4), the right prefrontal region (i.e., FFC6h), and the left motor cortex (i.e., CCP1h).

### Global structural and resting-state differences across stimulation conditions

With communities identified across stimulation conditions, we sought to determine the underlying causes for these differences. We hypothesized that the differences in TEP measures between the communities were driven by differences in the structural and resting state profile of the stimulation targets. The properties can be categorized as scalar or connectivity-based. Scalar measures include cortical thickness and myelination, which reflect the cortex’s properties at the stimulation target. They were calculated by averaging across the vertex-level values within each stimulation ROI.

Connectivity measures include those for DWI structural connectivity (SC), rsfMRI functional connectivity (RSFC), and rsEEG power envelope connectivity. For rsEEG, power envelope connectivity (PEC) was calculated for band-pass filtered data across the 4 canonical EEG frequency bands (theta, alpha, beta, gamma) separately for the resting eyes open (REO) and resting eyes closed (REC) conditions.

We chose degree as the summary metric for each connectivity measure, which was defined as the sum of the connectivity value between the E-field ROI and Schaefer 100 7-network parcels. A global summary metric was obtained by summing the connectivity values across all the parcels from a given stimulation ROI. Comparing the global structural and resting state measures across the communities showed significant group differences across multiple measures (Figure 4).

**Figure 4.**
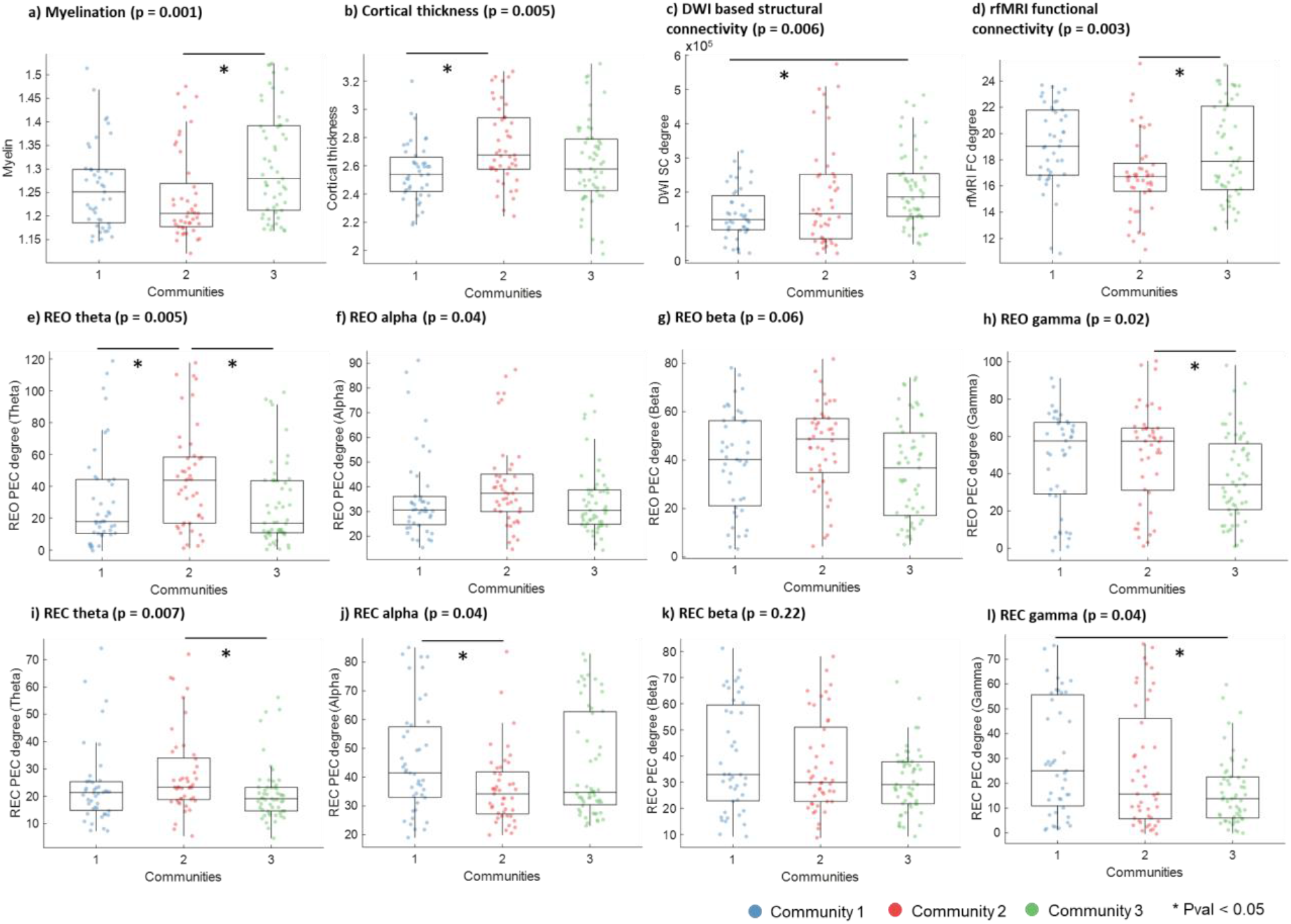
Comparing structural and resting state measures across the communities of the discovery set for measures of a) myelination, b) cortical thickness, c) DWI based structural connectivity, d) rfMRI functional connectivity, e) Stimulation area, f) SUD scale values. Overall group comparisons were done with Kruskal-Wallis test, while post-hoc paired comparisons between communities were done with a ranksum test. Significant paired comparisons after Tukey correction are marked by a star (Pval < 0.05).

Myelination was highest in community 3, significantly higher than in community 1. This aligns with previous results, as community 3 includes locations over the visual cortex (e.g., POz, Oz, MV1) and the motor cortex (e.g., C1, C2, CCP1h) that are shown to have high myelination (Glasser & Van Essen, 2011). SC revealed that the degree (or connectedness) of stimulation sites in community 3 was also highest. This suggests more white matter tracts to and from the stimulated regions for community 3. The fact that the community also has the highest cortical myelin bolsters this finding since the two are closely related (Glasser & Van Essen, 2011). Lastly, aligning with the DWI-based results, the RSFC also revealed that the degree or connectedness of stimulation sites in community 3 was highest, suggesting likely existence of hub regions in that community.

Differences in cortical thickness were also observed across communities, such that it was highest in community 2 and lowest in community 1. This result agrees with previous findings since the stimulation locations of community 2 are more concentrated in the frontal lobe, which is known for higher cortical thickness (Fischl & Dale, 2000).

Lastly, PEC measures were also different across communities. For REO PEC, community 2 has the highest value for all frequency bands. Given that community 2 has targets more concentrated in the frontal region, the increased theta and gamma PEC effects were expected (Toll et al., 2020). For REC PEC, community 2 has significantly higher theta than community 3, and significantly lower alpha than community 1. Comparing the frequency differences between REO and REC measures, one can see that while alpha PEC of community 2 is similar, there is an increase in the value for community 1, which is likely the driver of the effect. An increase in posterior and occipital alpha generators during the REC condition relative to REO condition is a known phenomenon, which supports our results here since community 2 has more targets in the frontal lobe and thus may be less affected by a boost in alpha synchrony.

### Network-specific structural and resting-state differences across stimulation conditions

In addition to comparing the global connectivity metrics between communities, resting state network-based summary metrics were obtained by dividing the sum of the connectivity values of parcels for each network by the total connectivity sum across all network parcels. Defined based on the Schaefer atlas with 7 networks and 100 parcels, the networks are default mode network (DMN), control network, limbic network, salient ventral attention network (VAN), dorsal attentional network (DAN), somatomotor network, and visual network (Schaefer et al., 2018). While the global metric quantifies the total connectivity from a stimulation ROI to the entire brain, the network-specific metric quantifies the connectivity to each network from a stimulation ROI relative to the total connectivity. Comparing the network-specific measures revealed additional insights into how each community differed in network connectivity distribution (**Supplemental Fig. S7 – S16**).

SC was significantly lower for the VAN network and higher for the DAN network in community 1 compared to community 2 (**Supplemental Fig. S7**), suggesting a reciprocal relationship between the two communities regarding structural connections.

RSFC was significantly higher for the VAN network but lower for the visual network in community 2 compared to community 3 (**Supplemental Fig. S8**), suggesting a reciprocal relationship between the two communities regarding functional connections.

REO theta PEC was significantly higher for the DMN and limbic networks but lower for the DAN and visual networks in community 2 compared to community 3 (**Supplemental Fig. S9**). Community 1 was also significantly higher in REO theta PEC for the DAN network compared to community 2. Similar to the case for REO, REC theta PEC was significantly higher for the DMN and limbic networks and lower for the DAN and visual networks in community 2 compared to community 3 (**Supplemental Fig. S10**). Community 1 was also significantly lower in REC theta PEC for the VAN network than community 3 and significantly higher for the visual network than community 2.

REO alpha PEC was significantly higher for the DMN and limbic networks but lower for the DAN network in community 2 compared to community 3 (**Supplemental Fig. S11**). REC alpha PEC included all the significant contrasts found for REO alpha PEC. In addition, community 1 and 2 had significantly lower REC alpha PEC for the control network than community 3, while community 1 had significantly lower values for the VAN network than community 2 (**Supplemental Fig. S12**).

REO beta PEC was significantly lower for the control network and VAN networks but higher for the visual network in community 1 compared to community 2. Community 2 also had significantly higher REO beta PEC for the DMN network than community 3 (**Supplemental Fig. S13**). Similarly, REC beta PEC was significantly lower for the control and VAN network, but higher for the visual network in community 1 compared to community 2 (**Supplemental Fig. S14**). In contrast, community 1 has significantly lower REC beta PEC for the visual network compared to community 2.

REO gamma PEC was significantly lower for the DMN, limbic, and VAN networks but higher for the DAN and visual networks in community 1 compared to community 2 (**Supplemental Fig. S15**). Community 2 was also significantly higher in REO gamma PEC for the limbic network compared to community 3. In contrast, the only significant difference for REC gamma PEC was lower values in community 1 than 2 for the limbic network (**Supplemental Fig. S16**).

### Similarity across recording channels

Complementary to the question of how stimulation sites are related to each other across all recording channels is how recording channels are related to each other across stimulation sites/orientations. This recording perspective was evaluated by calculating the correlation between all available stimulation conditions for a given channel, then correlating that similarity matrix across channels to finally arrive at a channel-by-channel similarity matrix used as input for community detection. Further details are provided in the methods section (Supplemental **Fig. S3**).

Across recording channels, community detection based on TEP similarity across channels also revealed three communities in the discovery set (Figure 5a), which, when applied to the validation set, directly revealed almost identical communities (Figure 5b). The communities are regional, but each has two separate components, labelled as subcomponents A and B (Figure 5c). Examining the voltage potential time series associated with each subcomponent shows that they are almost inversely correlated across time (Figure 5d).

**Figure 5.**
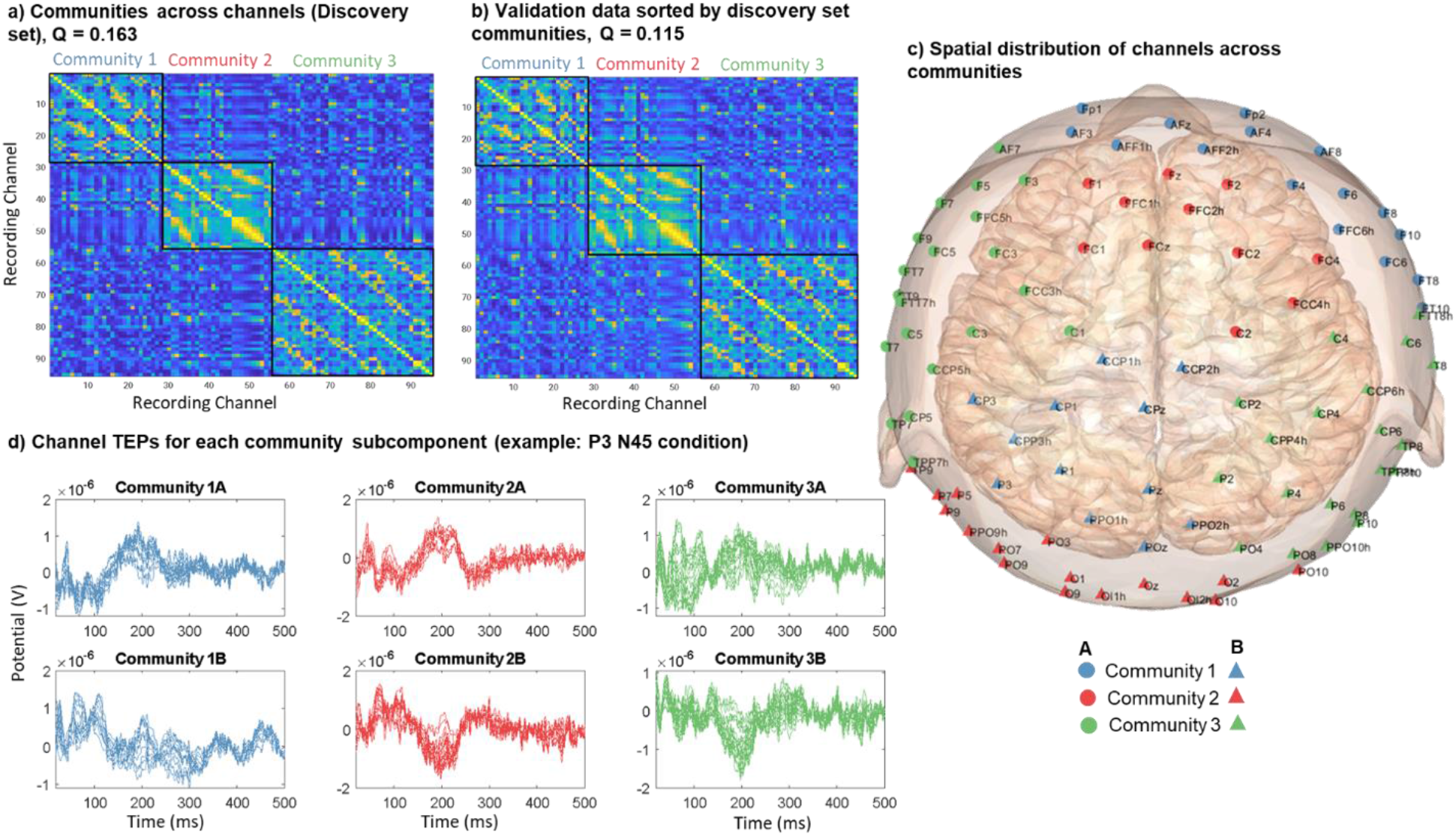
a) Community detection applied across recording channels based on TEP similarity in the discovery set, where Q is the modularity score. b) TEP similarity across channels for validation data sorted by the community labeling of the discovery set. c) Spatial distribution of channels across communities for the discovery set, with subcomponents labeled by different markers of the same color. d) Example of channel TEPs for each community subcomponent.

The channel community TEPs can be used to characterize each stimulation target as a dimensionality reduction method, which allows the TEP response to be summarized with 3 community time series (Supplemental Fig. S5 and S6). The same community time series can also be compared across conditions since they refer to the same channel groups. For example, community 2 has a large peak ∼ 150 ms for stimulation locations with orthogonal orientation over the motor cortex (N45 for the left hemisphere, P45 for the right hemisphere), while community 1 has a peak ∼ 115 ms over the CPz location for both coil orientations.

### Coil orientation sensitivity across recording channels

Similar to how different stimulation sites exhibited varying coil orientation sensitivity, different recording channels can also differ drastically in COS. To quantify COS for each recording channel, we calculated the modularity score of the TEP similarity matrix across all conditions using coil orientation as the community label. Before computing the correlation, conditions were split into orthogonal and parallel groups, which is in reference to the central sulcus for each hemisphere. A higher modularity score means a higher COS. To control for spurious findings, the actual score was compared against that of a null distribution. Further details are provided in the methods section (Supplemental Fig. S3).

For the discovery set, our results showed that channels with the highest modularity are localized to the left motor cortex (Figure 6a). When compared against the null distribution, the statistically significant channels (above 95% random values) are C3 and CCP5h. This means that across all the stimulation conditions, responses from those channels were most consistently different between the two coil orientations. For the validation set, our results showed additional channels with high modularity in addition to the left motor cortex, including ones in the frontal-central region and right occipital region (Figure 6b). The difference from the discovery set is likely due to the uneven distribution of stimulation conditions in the validation set. However, when compared with the null distribution specific to the validation set, the most significant channels are still over the left motor cortex, which includes C5 and CCP5h.

**Figure 6.**
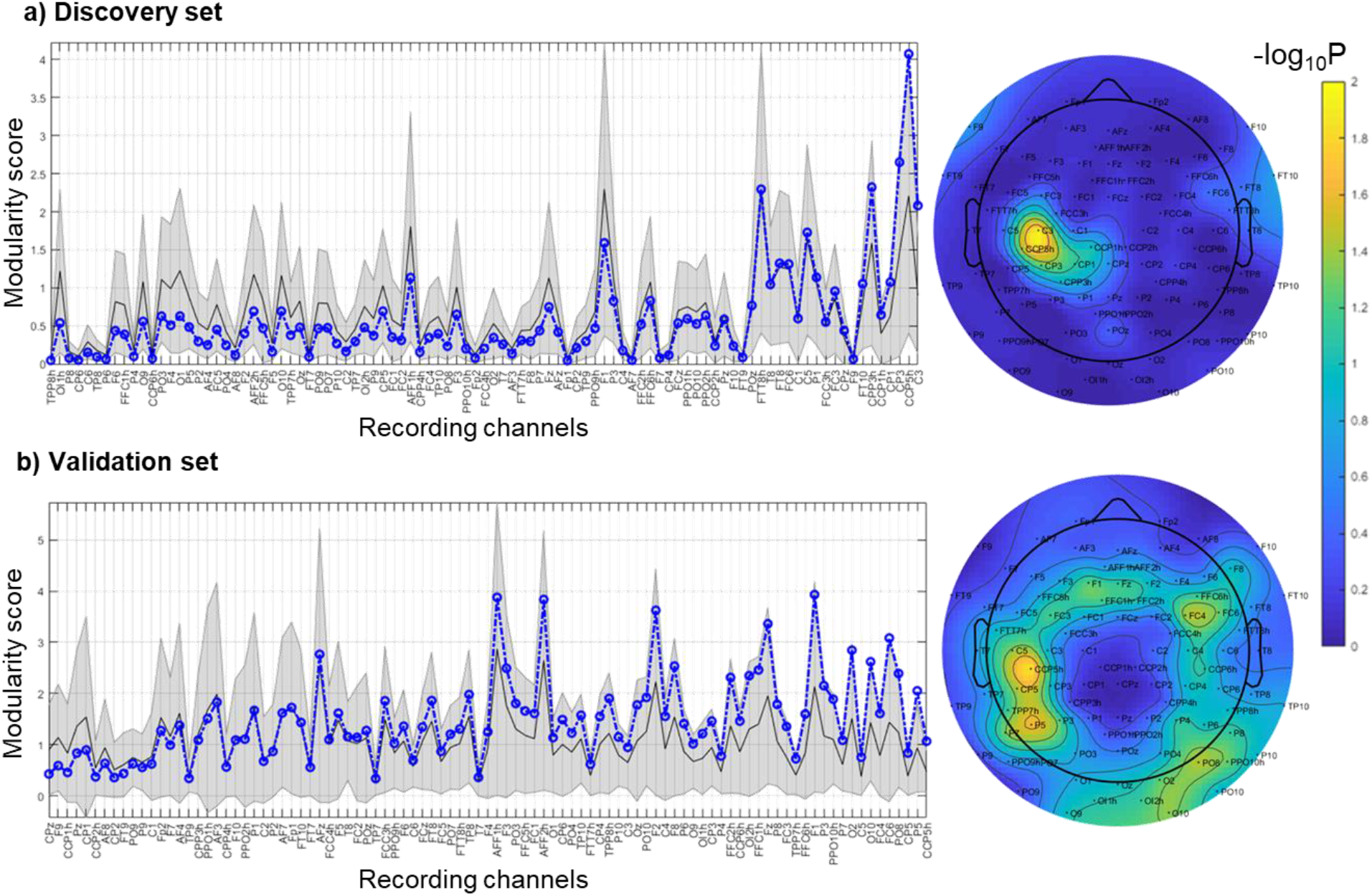
Significance of orientation modularity relative to null distribution for a) discovery set and b) validation set. The line plots show the observed modularity score (blue dashed line), along with the null distribution mean (black solid line) and 95% confidence intervals (grey shading). The values are sorted by ascending order of the significance of the observed value relative to null based on exact statistics (-log10P). The topoplots to the right show the significance values plotted spatially.

## Discussion

The release of this densely sampled N-of-1 dataset, both raw and processed, will provide a unique resource for both basic and applied neuroscientists to gain insights into the TMS-EEG stimulation response across cortical regions. Indeed, while TMS researchers may be familiar with the spatiotemporal characteristics of the TMS response for locations such as the motor cortex or prefrontal cortex, little is known about most other brain regions, including the timing of TEP peaks and their spatial distribution. This information can be visually examined quickly from the online lookup table or queried from the data, which is very useful for researchers hoping to study signals from a location part of the set or very close to one.

### Stimulation-based communities reveal distinct spatial and temporal patterns

Across all stimulation conditions, including both orientations, community detection with TEP connectivity identified 3 communities. The fact that only 3 distinct communities were identified suggests that from a network connectivity perspective, many stimulation conditions have similar effects. However, since the stimulation targets for each community were more interspersed rather than being localized spatially, they suggest underlying brain networks have complex borders and distributed components. These “TEP networks” may correspond to combinations of resting state networks.

Our results suggest that it may be possible to target different brain regions to achieve similar effects, which can, in turn, help future rTMS treatment studies to pick more accessible or tolerable targets for patients while maintaining similar treatment effects. At the same time, stimulation targets near the edge of the TEP networks may require extra attention as a small shift in positioning may lead to dramatically different effects, which concurs with previous literature (Chen et al., 2013).

Differences in the relative size of certain peaks in each community reflect differences in the network topology. For example, a larger early peak may reflect a shorter conduction time to a hub node in the network, which then activates many brain regions(Hens et al., 2019). More peaks for community 3 may suggest a network structure with more regional hubs.

Examination of the spatial distribution of the community TEPs revealed that certain prominent peaks are localized to specific scalp regions, which may represent specific brain processes. For example, the bilateral activation pattern for the 43 ms peak for community 1 is characteristic of the N45 component previously found in TEPs of several regions (Harquel et al., 2016; Kerwin et al., 2018; Lioumis et al., 2009). While the polarity of the N45 map is switched for targets of the two hemispheres, the absolute value average map shown in Figure 2 allows the process from both hemispheres to be treated the same. Likewise, the central midline pattern for the 193 ms component of community 2 is similar to the P200 component noted in previous TMS-EEG studies, while the midline frontal pattern for the 70 ms peak is similar to the P60 component (Harquel et al., 2016; Tremblay et al., 2019).

Based on previous literature, the N45 component has been associated with GABAa receptor-mediated neurotransmission (Premoli et al., 2014), the P60 with glutamatergic neurotransmission (Belardinelli et al., 2021), and the P200 with higher level information integration and long-range cortical connectivity (Massimini et al., 2005; Zipser et al., 2018). Since the N45, P60, and P200 components are visible for the TEPs of many targets (see online look-up table), the difference in the relative amplitude of the peaks between the communities may reflect the fact that they are still interconnected but provide different entry points and pathways into the same global network.

### Network-specific connectivity differences between stimulation-based communities

Our results showed that many structural and resting measures differed significantly between the communities, which may be used to delineate a neuroarchitecture profile for each community.

Stimulation sites of community 1 are characterized by higher REC alpha and gamma PEC but lower cortical thickness, SC, and REO theta PEC. Moreover, community 1 sites are associated with lower relative SC in the VAN network but higher relative SC in the DAN network. Across most of the frequency bands for both REO and REC, relative PEC connectivity was generally lower in the control, limbic, and VAN networks but higher in the DAN and visual networks. Therefore, together with the fact that the stimulation locations are more concentrated in the posterior lateral regions, this profile suggests that the accessed network may overlap with the DMN and DAN networks the most (Fox et al., 2006).

Stimulation sites of community 2 are characterized by higher cortical thickness, REO theta, and gamma PEC, and REC theta PEC, but lower myelination, RSFC, and REC alpha PEC. In terms of relative network SC, community 2 has higher values for the VAN network but lower values in the DAN network, which is flipped in comparison to community 1. In terms of relative network RSFC, community 2 has higher values for the VAN network but lower values for the visual network. Finally, in terms of PEC, community 2 is generally associated with higher values for the DMN, control, limbic, and VAN networks but lower values for the DAN and visual networks. Together with the fact that stimulation locations are more frontal central, this profile suggests that the accessed network may overlap with the VAN and control networks the most, which are more activated during wakefulness and operate in both theta and gamma frequency bands (Fries, 2015; Howard et al., 2003; Tallon-Baudry & Bertrand, 1999).

Finally, stimulation sites of community 3 are characterized by higher myelination, SC, and RSFC, but lower theta and gamma PEC for both REO and REC conditions. In terms of relative network RSFC, community 3 has higher values for the visual network but lower values for the VAN network, which is flipped in comparison to community 2. In terms of PEC, community 2 is generally associated with higher values for the DAN and visual networks but lower values for the DMN and limbic networks. Together with the concentration of stimulation sites over the occipital lobe, this profile suggests that the accessed network may overlap with the visual network the most, which is highly myelinated and connected by fast white matter tracts (Bullock et al., 2022; Glasser & Van Essen, 2011).

While these TMS stimulation-based communities may share common traits with known resting state networks, details of the causal network processes may be uniquely captured through TMS-EEG mapping. The profiles of the communities will help researchers determine where to stimulate in order to preferentially activate specific networks.

### Coil orientation sensitivity across stimulation locations

From a stimulation perspective, we found that most stimulation locations are sensitive to coil orientation. Only about a third of the targets were considered to have low COS based on community matching between the two coil angles. Those targets with low COS are more concentrated in the posterior central and occipital regions. Comparing the dissimilarity values qualitatively shows that regions near M1 (e.g., CCP1h, C4) have the highest COS, as expected based on previous studies (Bonato et al., 2006; Casula et al., 2022), but also the right frontal region (e.g., F4, FFC6h) was observed to have high COS.

Mechanistically, a stimulation location with high COS indicates anisotropy in the underlying brain structure. This is exemplified by the motor cortex, which is organized somatotopically along the pre-central gyrus. A stimulation location with low COS indicates structural isotropy, which is more likely in brain regions with more complex and less regular folding patterns, such as the posterior cortex.

Practically, a region with high COS requires more attention when positioning the coil. For example, maintaining the same angle between conditions and subjects may be necessary to reduce unwanted variability for comparisons. Alternatively, two or more coil orientations may be employed during data collection to capture a more unbiased average response from the region.

It is worth noting that the number of communities matters when examining community agreements. The more communities are identified, the less likely the two paired conditions will match. For example, if there were 2 equal communities randomly distributed for each orientation, then the expected number of matches by chance would be 1/2. Similarly, for 3 equal size communities, the expected number of matches by chance would be 1/3.

### Characteristic regional responses irrespective of stimulation condition

Communities identified across recording channels show there are characteristic responses of regions irrespective of stimulation condition. This partially arises because of the fixed structural connectome. Regardless of where the initial stimulus is, the response will involve the same structural components, albeit at different levels and timings. The presence of two subcomponents to each identified community points to dipole-like sources that might underlie the observed sensor-level activity. It also illustrates how current flows as part of brain processes are conserved within the brain, where net inflow to one region requires a net outflow from another region.

In terms of practical application, having the same channel communities for all stimulation sites provides a low-dimensional view of the TMS-EEG responses, which may assist with identifying patterns across the conditions. Moreover, the communities’ spatial topography can help design lower-density EEG systems for clinical applications. For example, by placing one channel at the centroid of each community (or community subcomponent), researchers can putatively maximize the amount of information gathered with the few recording channels, which is especially helpful in a clinical setting where faster setup is imperative.

### Coil orientation sensitivity across recording channels

From a recording perspective, channels over the left motor cortex have the highest COS in the discovery and validation set. This finding may be due to the high orientation sensitivity of the left motor cortex when activated, which may cause it to be more sensitive to changes in input signals from other activated brain regions (Rizzolatti & Luppino, 2001). Since the participant was right-handed, the dominant left motor cortex is likely more represented and connected in the brain (Nicolini et al., 2019).

A potential application of quantifying COS from a recording perspective is optimizing treatments for modulating a distal target. Indeed, the relationship between a cortical target and a deeper structure (e.g., anterior cingulate) can be established by applying source localization techniques. When the COS is high for such a recording region, the need for coil adjustments may be more critical.

### Limitations and analysis prospects

A potential concern for all TMS-EEG studies is that auditory and somatosensory evoked potentials may still affect the preprocessed TEP signal (Conde et al., 2019). We have taken experimental precautions (i.e., auditory masking, foam padding the coil) to minimize potential effects. We also regressed out the effect of SUDs before all our analyses.

An inherent limitation for a N-of-1 study is the question of generalizability. However, using deeply phenotyped individual results has helped advance the field of neuroscience in understanding common population traits. Such an approach is common for non-human primate experiments where long and involved behavioral experiments are performed with one or few subjects (Wessberg et al., 2000). Similar approaches have been used for establishing the Talairach coordinate system (Talairach & Tournoux, 1988) for creating the Colin27 template MRI brain (Holmes et al., 1998). Nevertheless, a future extension of our work is to create a more generalizable population representative stimulation-response map based on a larger cohort of subjects.

A sparser grid will be required since running more than 200 conditions per person is not realistically scalable. However, an immediate challenge is deciding which sparse targets to include. For connectome mapping, targeting each of the TEP-based communities can be a heuristic for generating an efficient sparse map. One target from each community can be sampled first before additional targets from the same community are obtained to maximize information gathering. In this sense, the present N-of-1 study provided a first sketch of a common landscape depicting the stimulation-response profile across the human cortex, making future efforts to create a population template more tractable.

## Conclusion

With over 100 stimulation locations and two coil orientations each, this study has the highest density of recorded TMS-EEG responses across the brain in one individual. An online lookup table was created for researchers to quickly check each stimulation condition’s response profile before further examining the data. For researchers hoping to get an overview of the stimulation-response characteristics across the brain, our study showed that community detection applied to TEP measures could reveal insights about the brain’s organization, both from a stimulation and recording perspective. Our results also showed that structural and resting measures differed significantly between TMS-EEG-based communities identified across stimulation conditions, revealing differences in the neuroarchitecture profile of each community. Given the availability of high-quality whole-brain neuroimaging data in addition to the TMS-EEG recordings, this neuro resource is perfect for building and validating models of TMS-EEG response that depend on the underlying structural and resting state measures. Likewise, the resource may be developed into a teaching example for demonstrating how TMS-EEG responses relate to other neuroimaging modalities.

## Data and code availability

The raw and processed data will be shared on OpenNeuro upon publication. The TMS-EEG preprocessing pipeline is maintained in an online repository (https://github.com/chriscline/AARATEPPipeline). Any additional information required to reanalyze the data reported in this paper is available from the lead contacts upon request.

## Author contributions

Conceptualization: YS, MVL, AE, MS

Investigation: YS, MVL, MM, MS

Methodology: YS, MVL, CCC, SK, FB, MN, WW, MS

Analyzed Data: YS, MVL, CCC, SK, MS

Writing—Original Draft: YS, MVL, AE, MS

Writing—Review & Editing: YS, MVL, CCC, MM, SK, FB, MN, WW, ZJD, AE, MS

Visualization: YS, CCC

## Acknowledgements

This work was supported by a NIH Director’s New Innovation Award (MH119735) and a NIMH R01 (MH127608) to M.S., and a NIH Director’s Pioneer Award (MH116506) to A.E. Funding support was also provided through a Canadian Institutes of Health Research (CIHR) Fellowship (Y.S.), a Stanford Bio-X Fellowship (M.V.L.), and a Department of Veterans Affairs Mental Illness Research, Education, and Clinical Center (MIRECC) Advanced Fellowship (C.C.C.). The authors thank Russell A. Poldrack for his guidance in approaching N of 1 studies.

