## Supplemental information for "Densely sampled stimulus-response map of human cortex with single pulse TMS-EEG and its relation to whole brain neuroimaging measures"

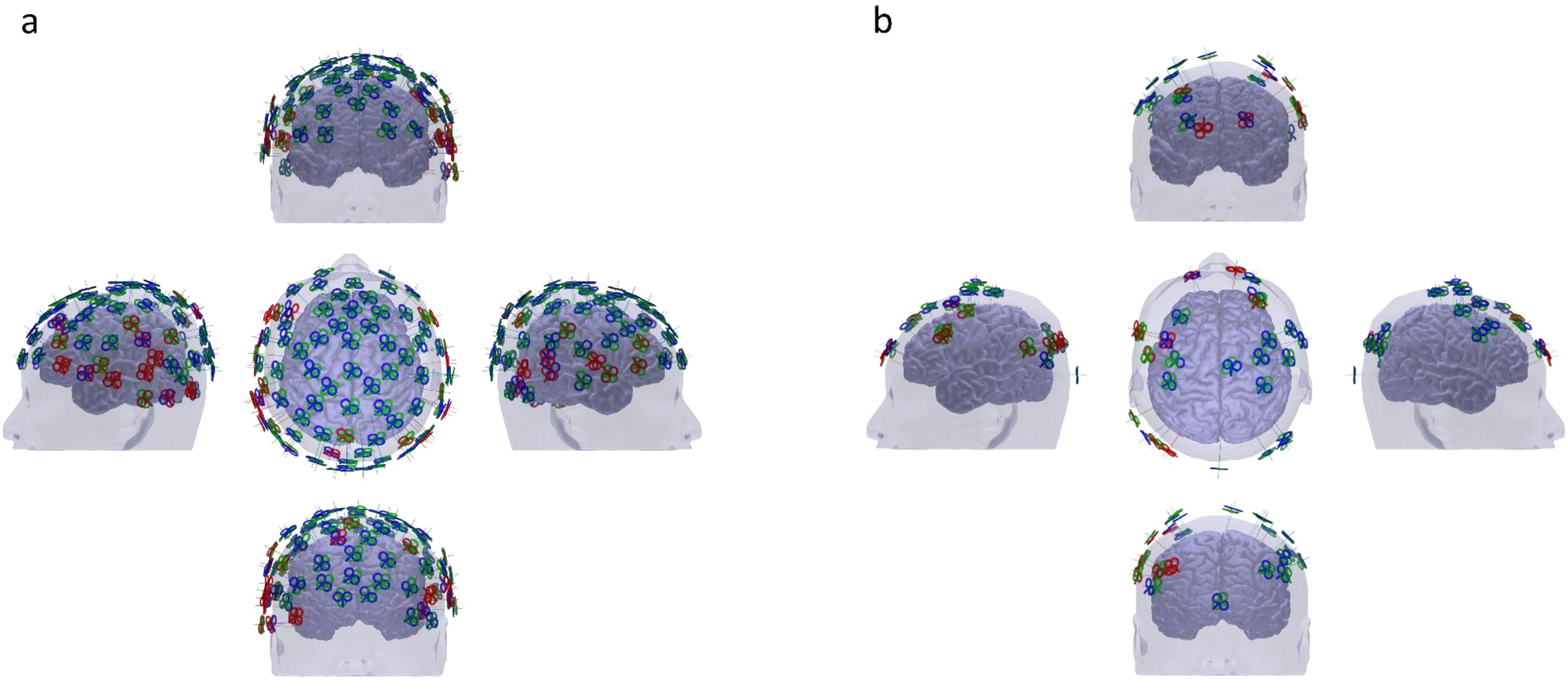


***Supplemental Fig. S1.*** *Quality control results of TMS-EEG data across (a) main targets, and (b) validation targets. Target coils labeled in red were not included in final analysis.*

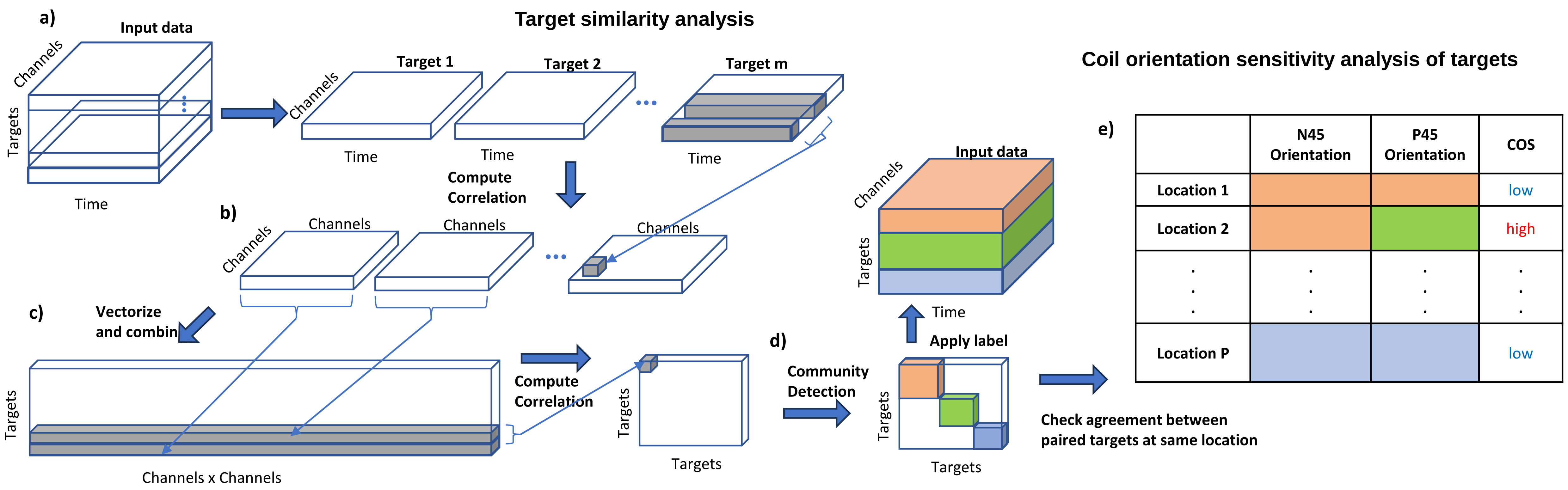


***Supplemental Fig. S2.*** *For each stimulation target, the TEP data matrix (95 channels x 481 timepoints, panel a) was correlated across timepoints to obtain a 95 x 95 channel connectivity matrix (panel b), which was then reshaped into a 1 x 9025 vector and combined (panel c). The resulting targets by channel connectivity matrix was correlated across all connectivity values to produce a targets x targets matrix ($A_{ij}$ in equation 1) that was used as input for community detection (panel d). Coil orientation sensitivity (COS) for each stimulation location was quantified by the community agreement between the associated P45 and N45 targets (panel e). If paired targets were assigned different communities, then the shared location was considered to have high COS.*

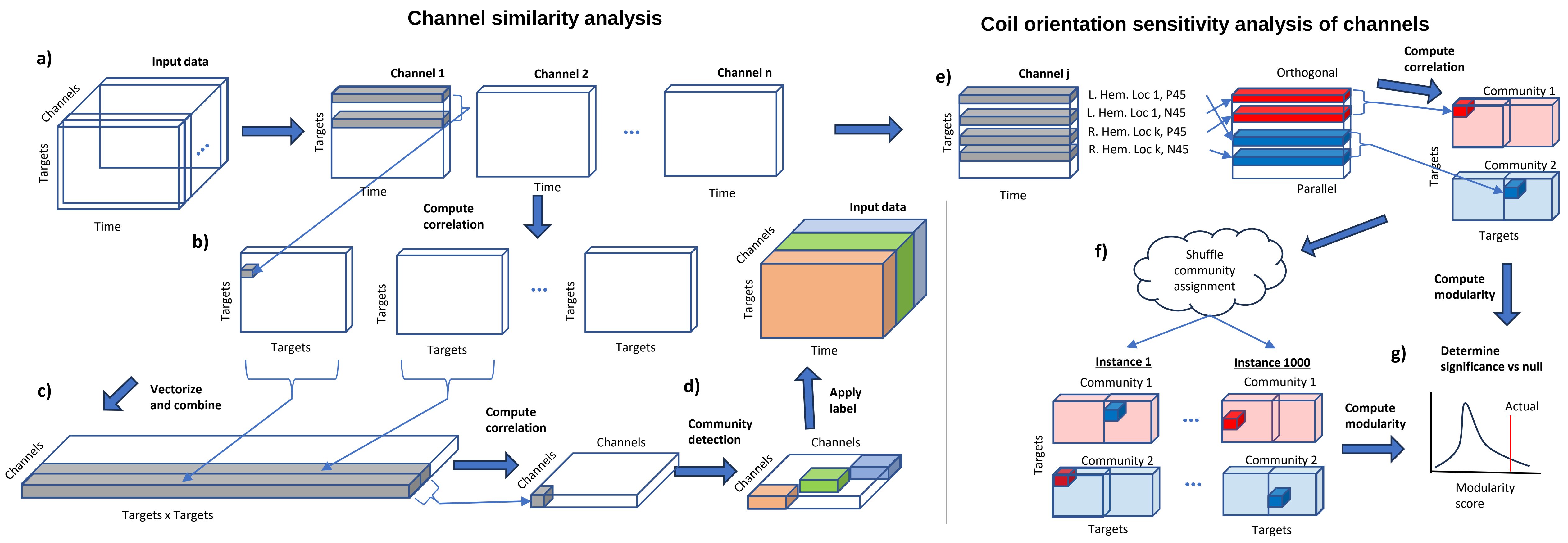


***Supplemental Fig. S3.*** *For each recording channel, an associated targets x timepoints matrix (panel a) can be correlated across timepoints to obtain a targets x targets similarity matrix (panel b), which can then be reshaped into a vector and combined. The resulting channels by targets connectivity matrix (panel c) was used as the input ($A_{ij}$) for community detection (panel d). COS for each channel $j$ was quantified by the modularity score of the TEP timeseries across all targets using coil orientation as the community label (panel e). Targets were split into orthogonal and parallel groups, which is in reference to the central sulcus for each hemisphere, before computing the correlation. A higher modularity score means a higher COS. A null distribution was generated by shuffling the community assignments and then computing the modularity score by chance for 1000 times (panel f). A significance test ($p$ < 0.05, or above 95% random values) was performed by comparing the actual modularity against that of the null distribution using exact statistics (panel g).*

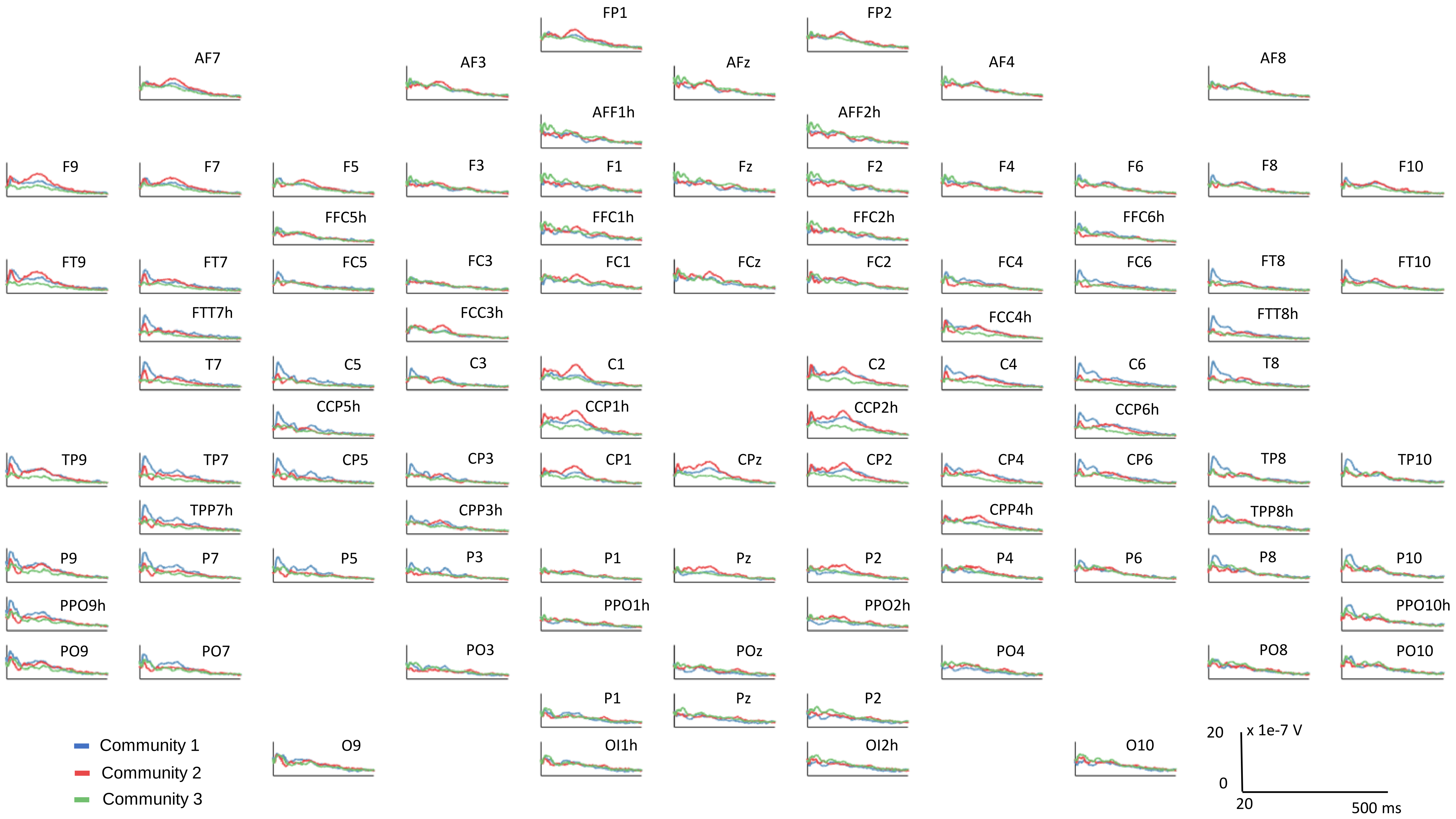


***Supplemental Fig. S4.*** *Channel TEPs averaged across targets of same communities (Discovery set, average of absolute value)*

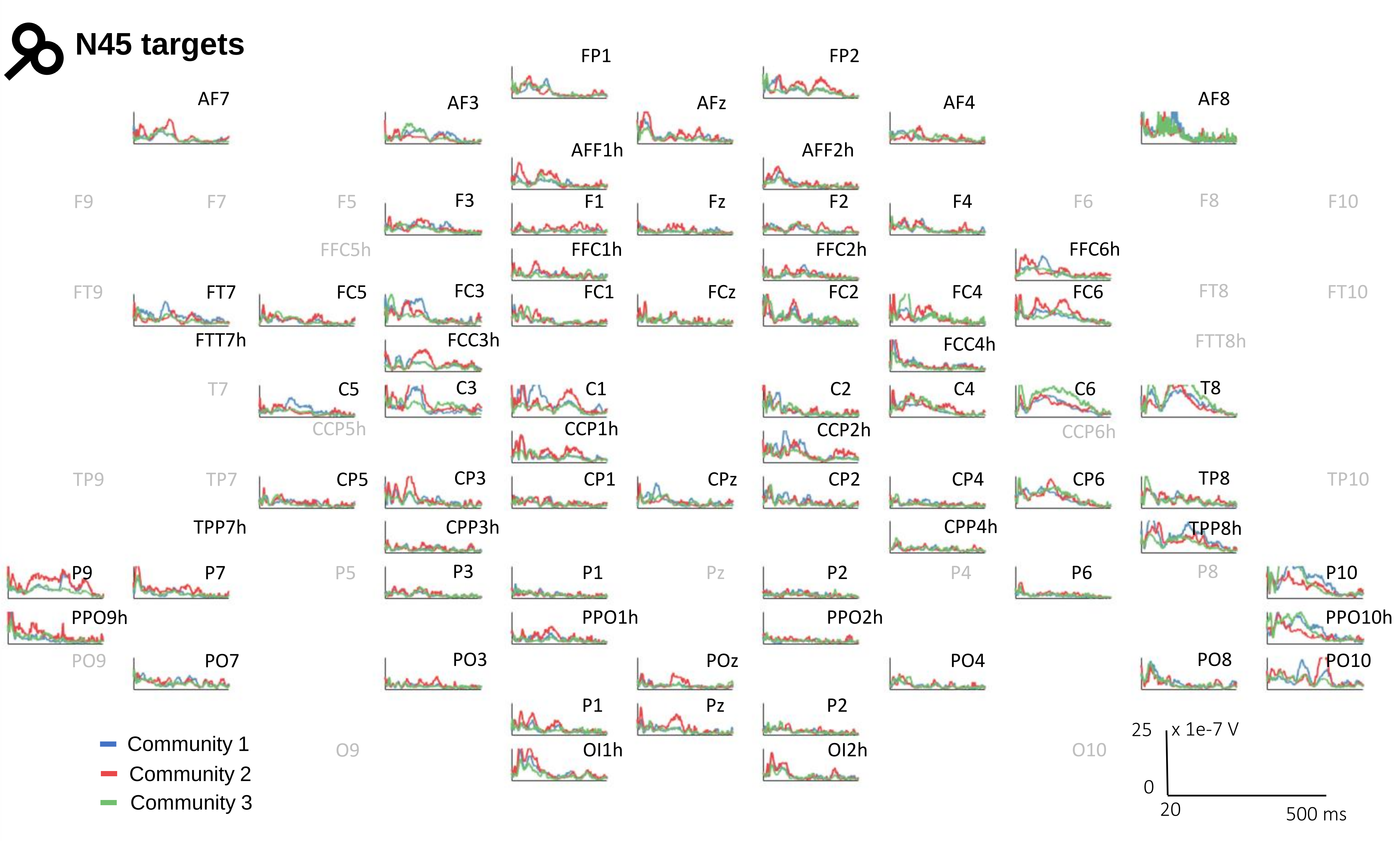


***Supplemental Fig. S5.*** *Community timeseries (absolute value average) for each N45 target of discovery set. Each electrode position corresponds to a stimulation target at that location. The greyed-out ones were either skipped due to experiment issues or did not pass the quality control check.*

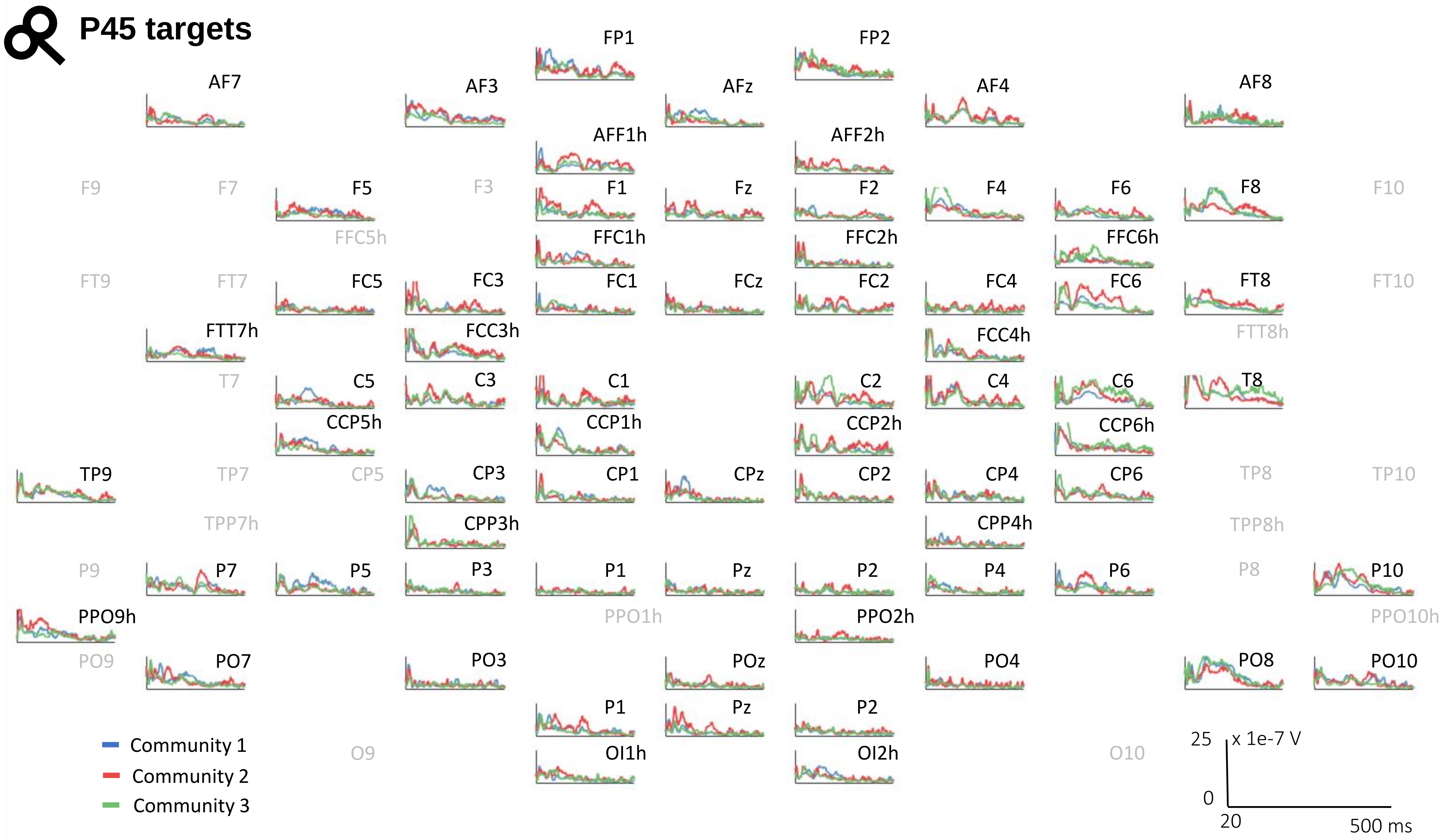


***Supplemental Fig. S6.*** *Community timeseries (absolute value average) for each P45 target of discovery set. Each electrode position corresponds to a stimulation target at that location. The greyed-out ones were either skipped due to experiment issues or did not pass the quality control check.*

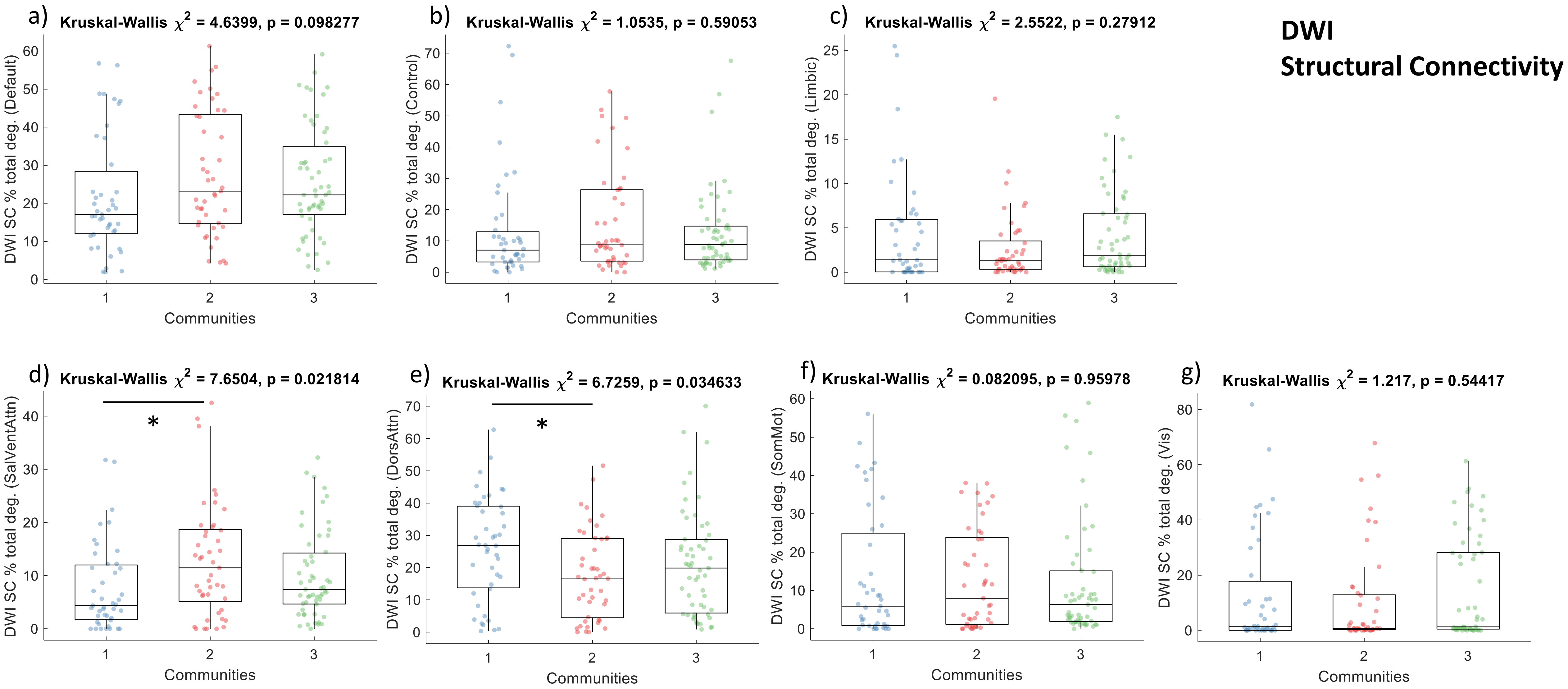


***Supplemental Fig. S7. Comparing DWI structural connectivity (SC)*** *across the communities of the discovery set based on network-specific degree as a percentage of the total degree, for the a) default mode, b) control, c) limbic, d) salient ventral attention, e) dorsal attention, f) somatomotor, and g) visual networks. Overall group comparisons were done with Kruskal-Wallis test, while post-hoc paired comparisons between communities were done with a ranksum test. Significant paired comparisons after Tukey correction are marked by a star (Pval < 0.05).*

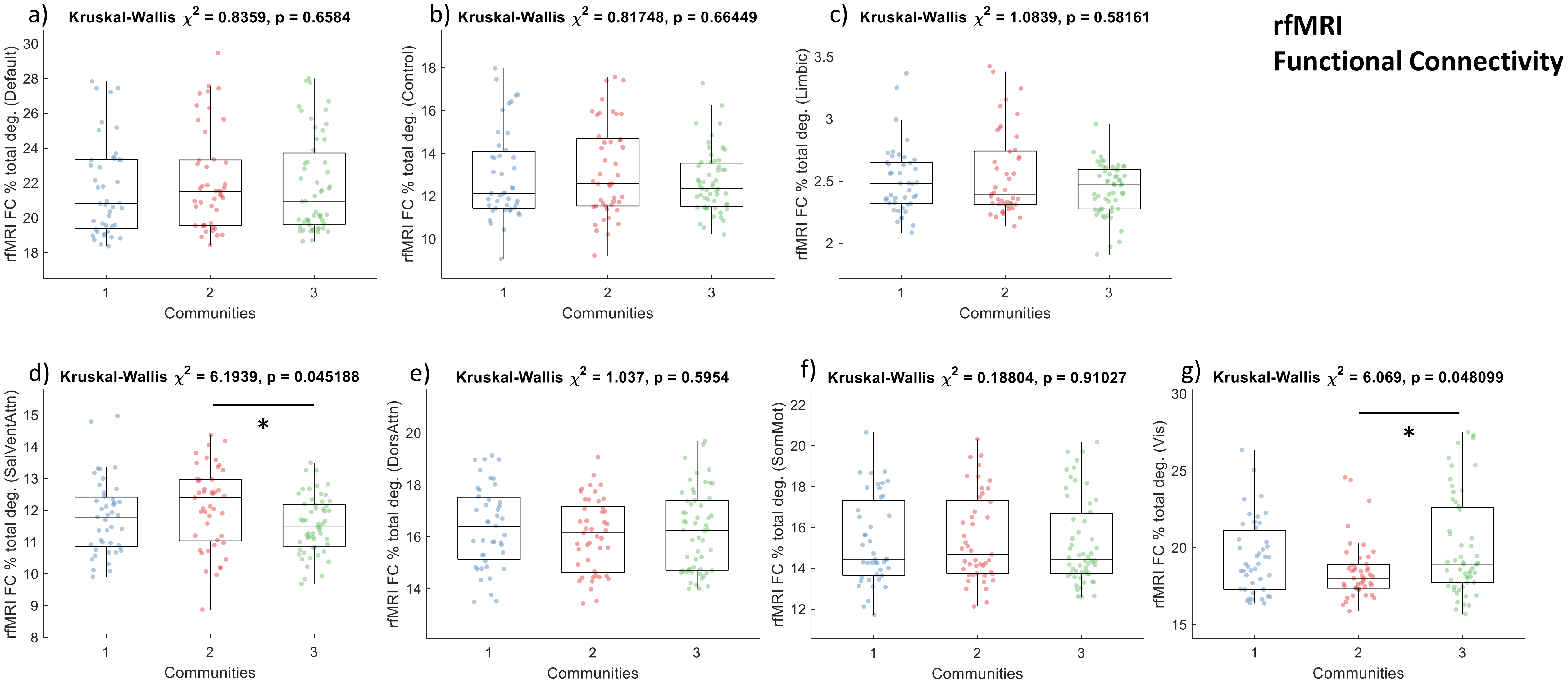


***Supplemental Fig. S8. Comparing rfMRI functional connectivity (FC)*** *across the communities of the discovery set on network-specific degree as a percentage of the total degree, for the a) default mode, b) control, c) limbic, d) salient ventral attention, e) dorsal attention, f) somatomotor, and g) visual networks. Overall group comparisons were done with Kruskal-Wallis test, while post-hoc paired comparisons between communities were done with a ranksum test. Significant paired comparisons after Tukey correction are marked by a star (Pval < 0.05).*

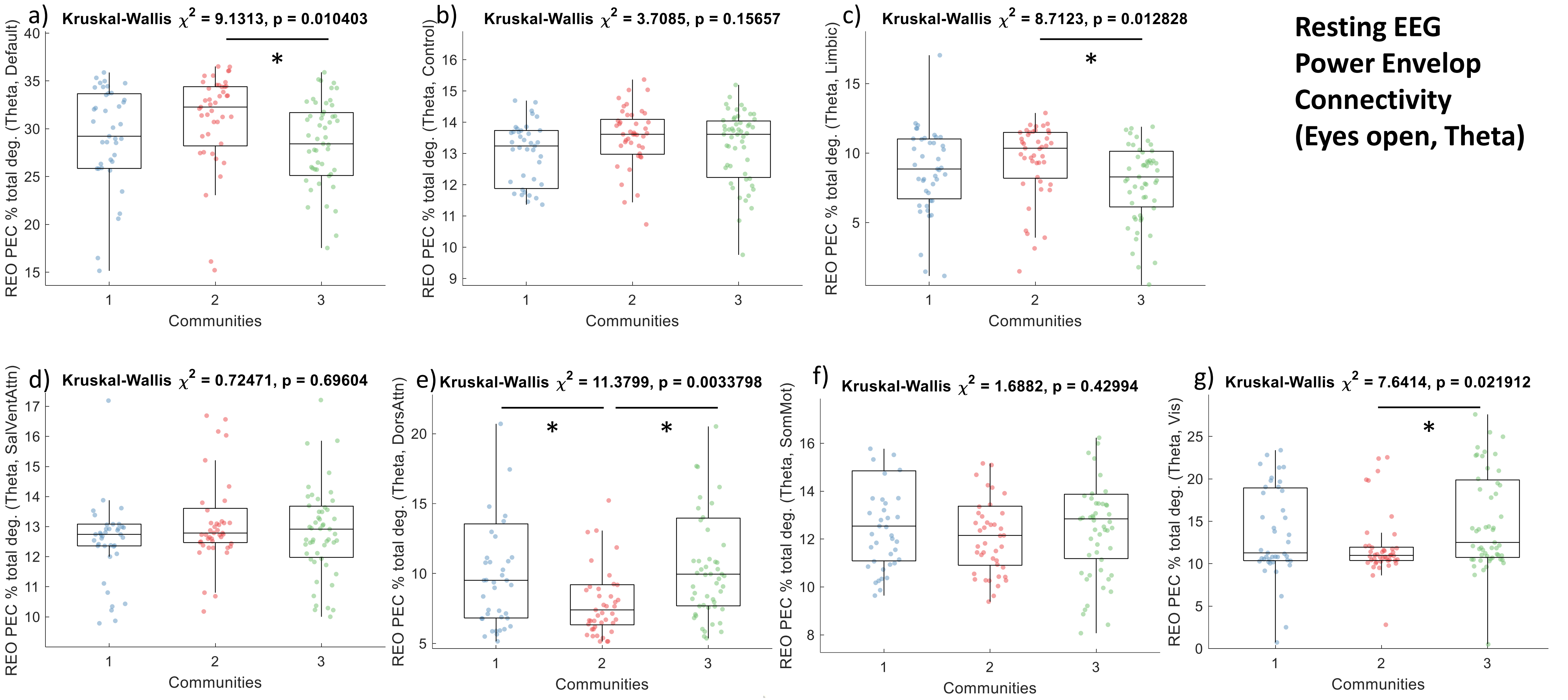


***Supplemental Fig. S9. Comparing resting eyes open (REO) power envelop connectivity (PEC) for the theta band*** *across the communities of the discovery set on network-specific degree as a percentage of the total degree, for the a) default mode, b) control, c) limbic, d) salient ventral attention, e) dorsal attention, f) somatomotor, and g) visual networks. Overall group comparisons were done with Kruskal-Wallis test, while post-hoc paired comparisons between communities were done with a ranksum test. Significant paired comparisons after Tukey correction are marked by a star (Pval < 0.05).*

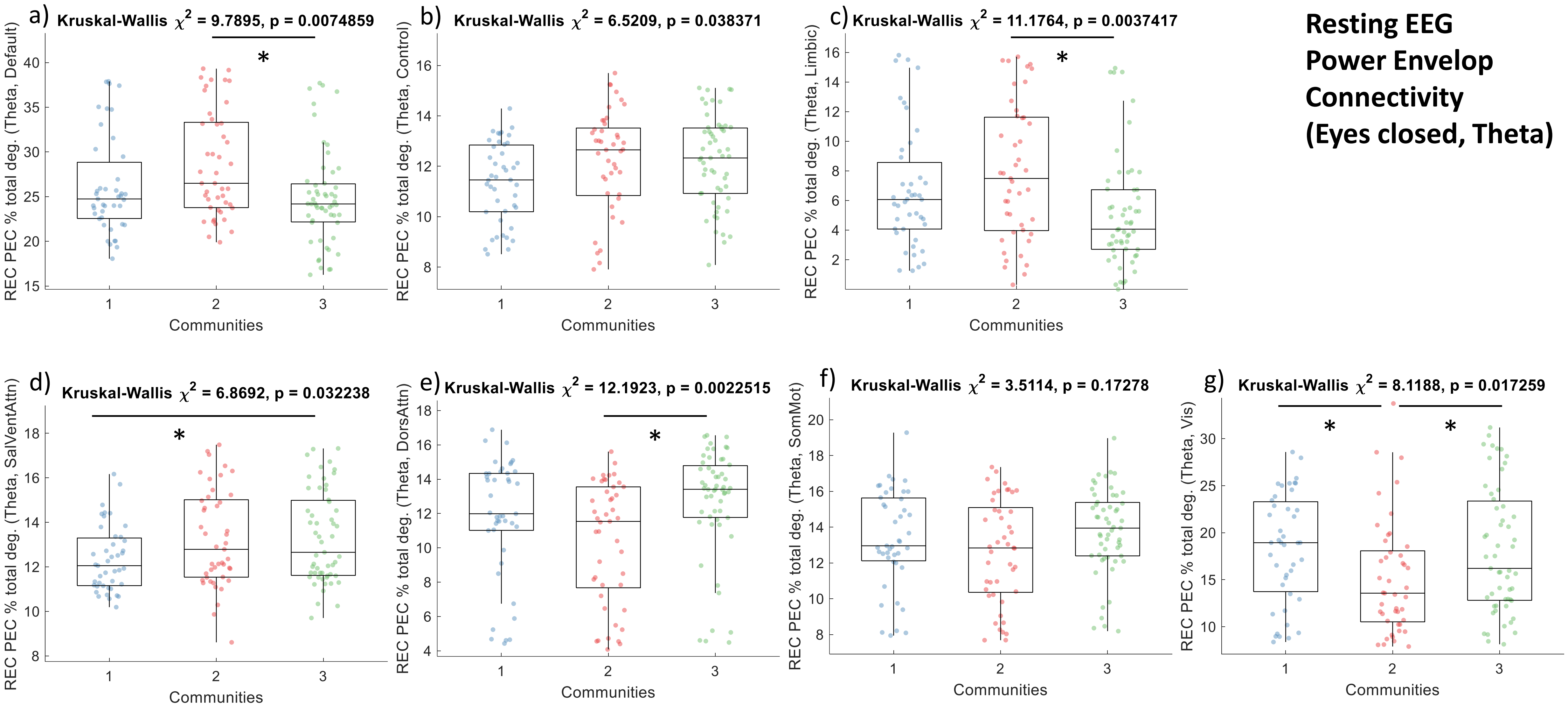


***Supplemental Fig. S10. Comparing resting eyes closed (REC) power envelop connectivity (PEC) for the theta band*** *across the communities of the discovery set on network-specific degree as a percentage of the total degree, for the a) default mode, b) control, c) limbic, d) salient ventral attention, e) dorsal attention, f) somatomotor, and g) visual networks. Overall group comparisons were done with Kruskal-Wallis test, while post-hoc paired comparisons between communities were done with a ranksum test. Significant paired comparisons after Tukey correction are marked by a star (Pval < 0.05).*

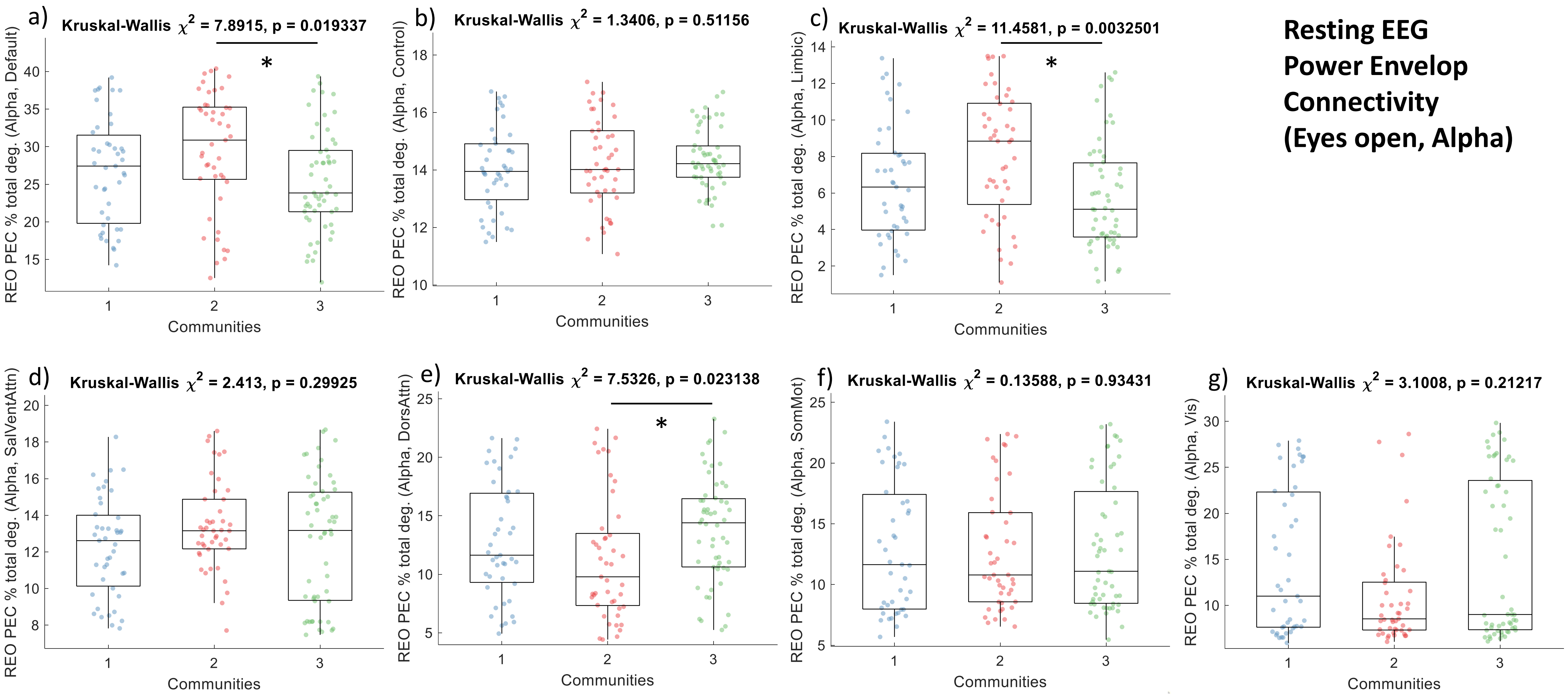


***Supplemental Fig. S11. Comparing resting eyes open (REO) power envelop connectivity (PEC) for the alpha band*** *across the communities of the discovery set on network-specific degree as a percentage of the total degree, for the a) default mode, b) control, c) limbic, d) salient ventral attention, e) dorsal attention, f) somatomotor, and g) visual networks. Overall group comparisons were done with Kruskal-Wallis test, while post-hoc paired comparisons between communities were done with a ranksum test. Significant paired comparisons after Tukey correction are marked by a star (Pval < 0.05).*

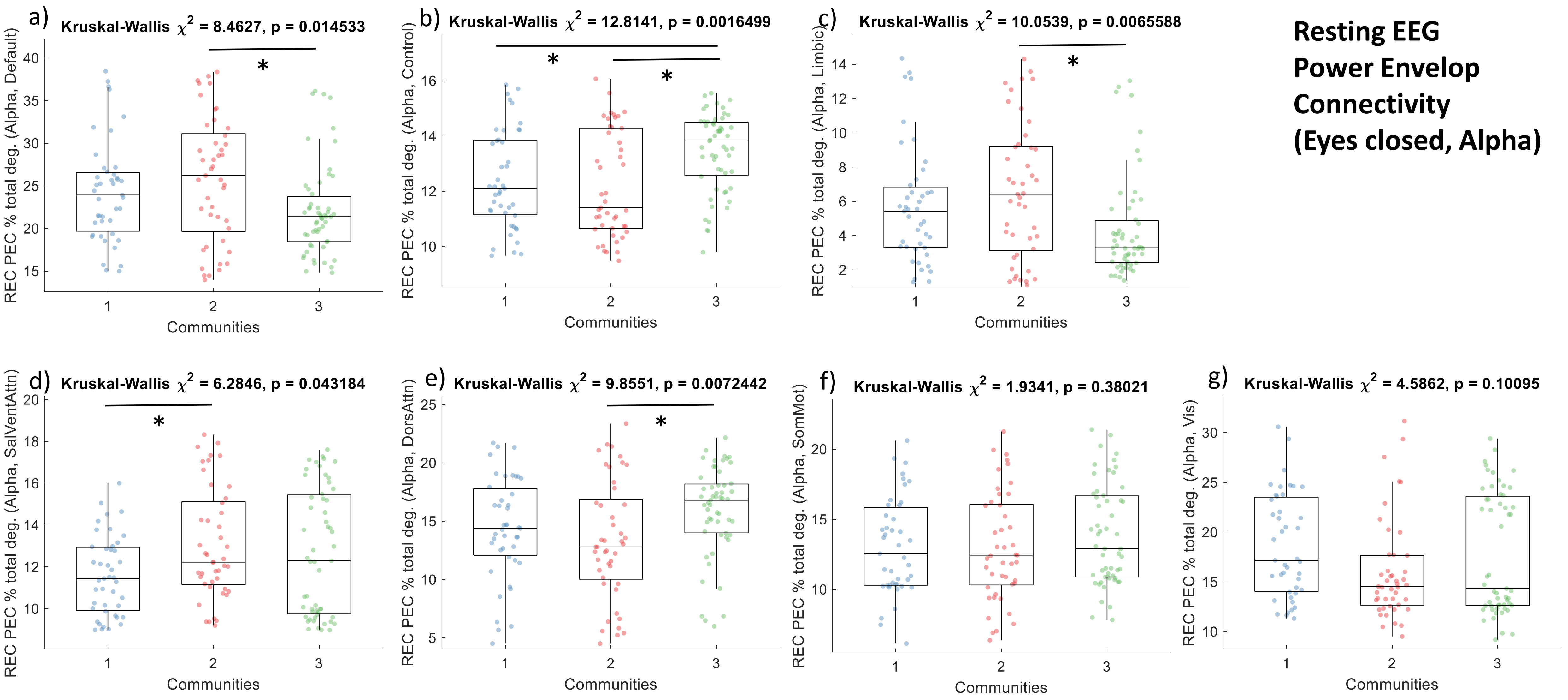


***Supplemental Fig. S12. Comparing resting eyes closed (REC) power envelop connectivity (PEC) for the alpha band*** *across the communities of the discovery set on network-specific degree as a percentage of the total degree, for the a) default mode, b) control, c) limbic, d) salient ventral attention, e) dorsal attention, f) somatomotor, and g) visual networks. Overall group comparisons were done with Kruskal-Wallis test, while post-hoc paired comparisons between communities were done with a ranksum test. Significant paired comparisons after Tukey correction are marked by a star (Pval < 0.05).*

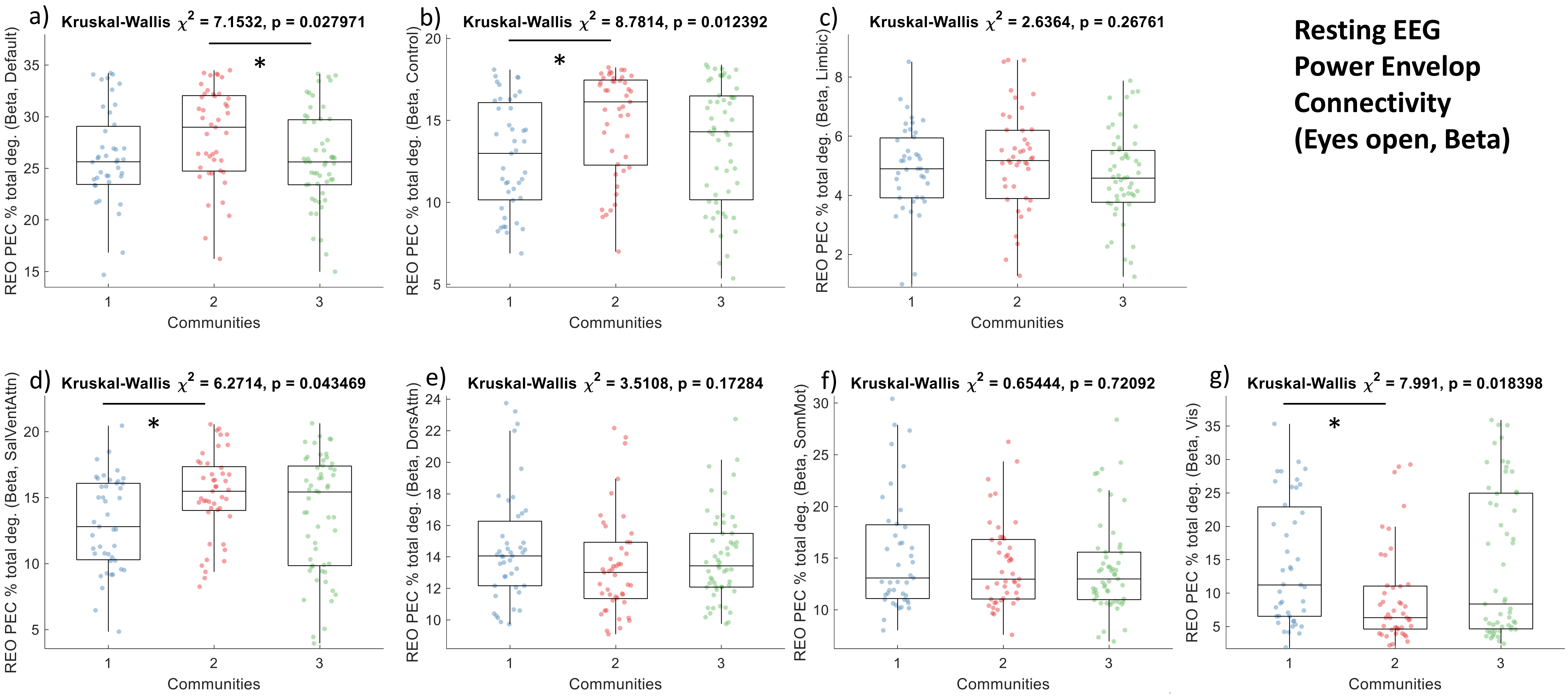


**Resting EEG Power Envelop Connectivity (Eyes open, Beta)**

***Supplemental Fig. S13. Comparing resting eyes open (REO) power envelop connectivity (PEC) for the beta band*** *across the communities of the discovery set on network-specific degree as a percentage of the total degree, for the a) default mode, b) control, c) limbic, d) salient ventral attention, e) dorsal attention, f) somatomotor, and g) visual networks. Overall group comparisons were done with Kruskal-Wallis test, while post-hoc paired comparisons between communities were done with a ranksum test. Significant paired comparisons after Tukey correction are marked by a star (Pval < 0.05).*

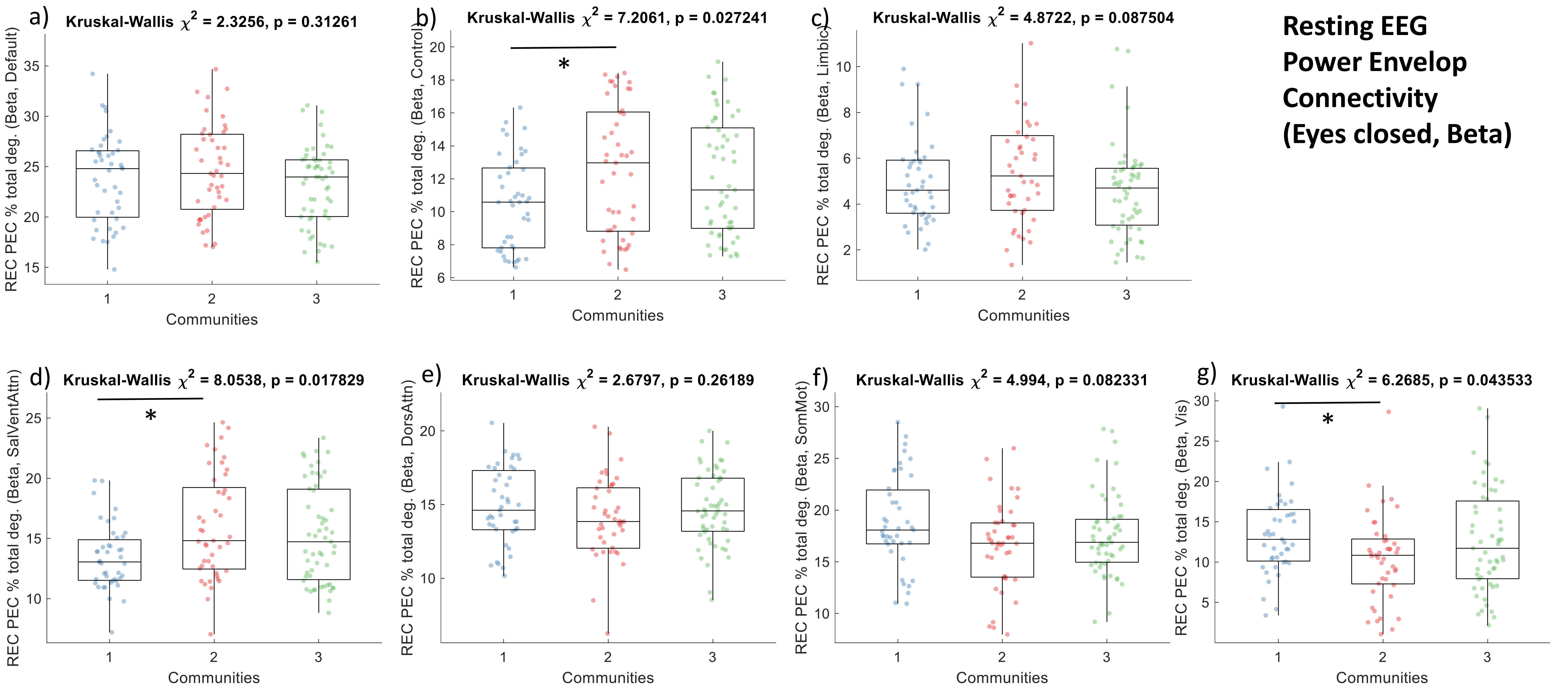


***Supplemental Fig. S14. Comparing resting eyes closed (REC) power envelop connectivity (PEC) for the beta band** across the communities of the discovery set on network-specific degree as a percentage of the total degree, for the a) default mode, b) control, c) limbic, d) salient ventral attention, e) dorsal attention, f) somatomotor, and g) visual networks. Overall group comparisons were done with Kruskal-Wallis test, while post-hoc paired comparisons between communities were done with a ranksum test. Significant paired comparisons after Tukey correction are marked by a star (Pval < 0.05).*

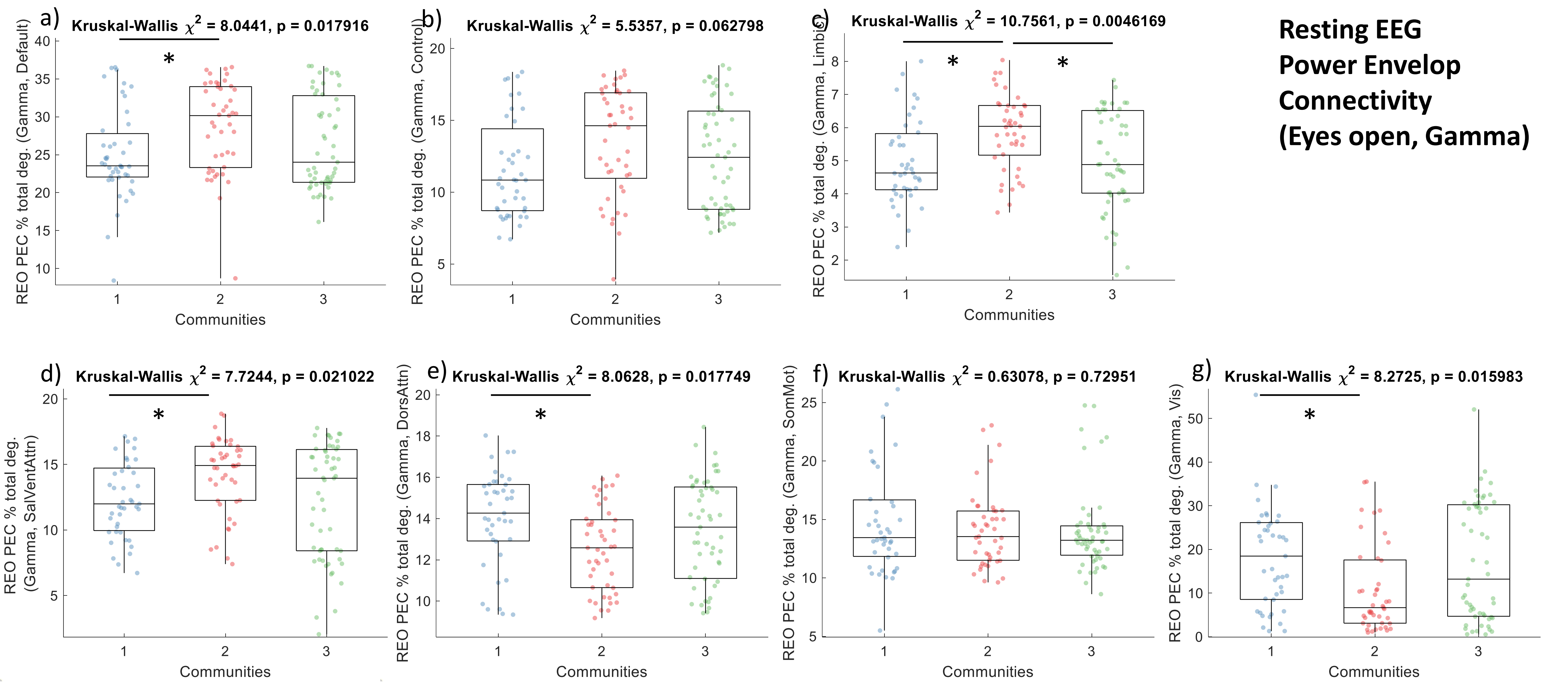


***Supplemental Fig. S15. Comparing resting eyes open (REO) power envelop connectivity (PEC) for the gamma band*** *across the communities of the discovery set on network-specific degree as a percentage of the total degree, for the a) default mode, b) control, c) limbic, d) salient ventral attention, e) dorsal attention, f) somatomotor, and g) visual networks. Overall group comparisons were done with Kruskal-Wallis test, while post-hoc paired comparisons between communities were done with a ranksum test. Significant paired comparisons after Tukey correction are marked by a star (Pval < 0.05).*

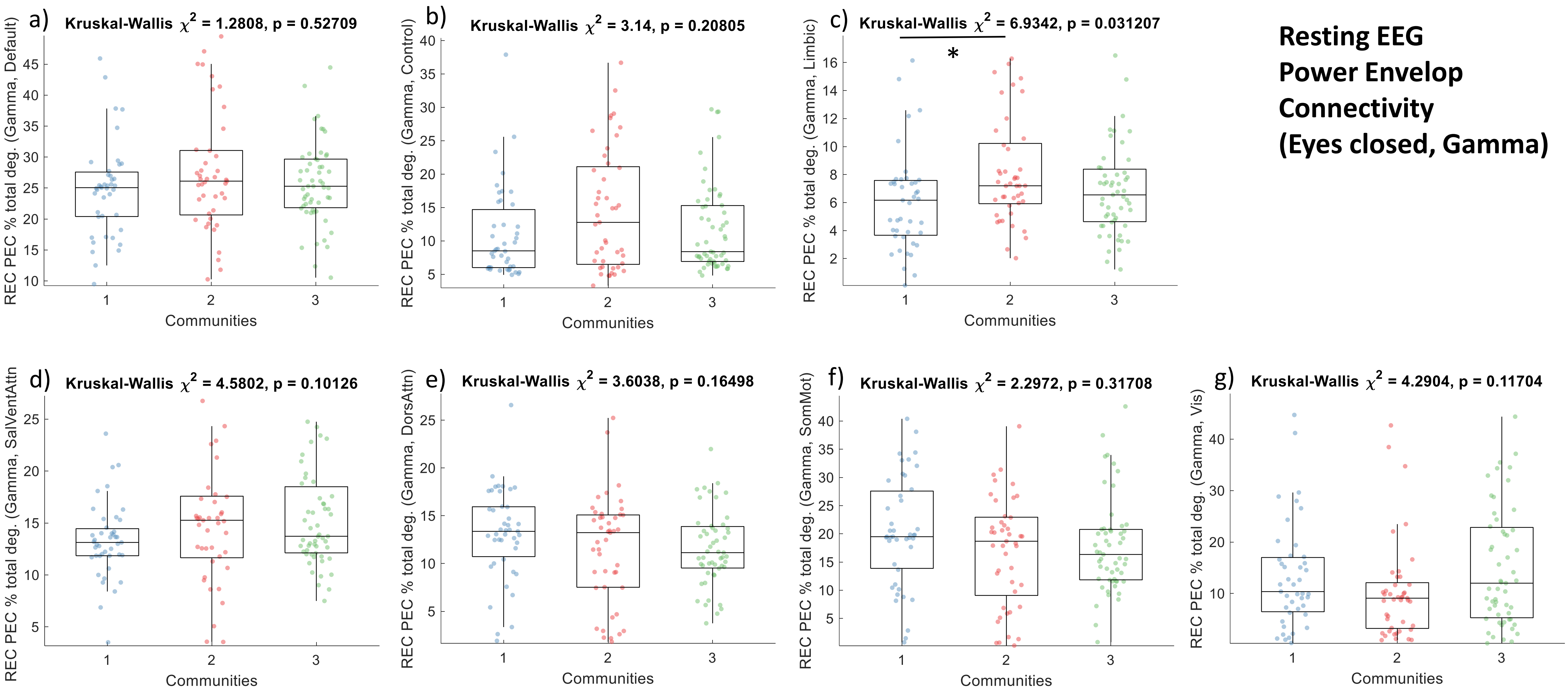


***Supplemental Fig. S16. Comparing resting eyes closed (REC) power envelop connectivity (PEC) for the gamma band*** *across the communities of the discovery set on network-specific degree as a percentage of the total degree, for the a) default mode, b) control, c) limbic, d) salient ventral attention, e) dorsal attention, f) somatomotor, and g) visual networks. Overall group comparisons were done with Kruskal-Wallis test, while post-hoc paired comparisons between communities were done with a ranksum test. Significant paired comparisons after Tukey correction are marked by a star (Pval < 0.05).*
